# Subfamily evolution analysis using nuclear and chloroplast data from the same reads

**DOI:** 10.1101/2024.01.15.575800

**Authors:** Eranga Pawani Witharana, Takaya Iwasaki, Myat Htoo San, Nadeeka U. Jayawardana, Nobuhiro Kotoda, Masashi Yamamoto, Yukio Nagano

## Abstract

The chloroplast (cp) genome is a widely used tool for exploring plant evolutionary relationships, yet its effectiveness in fully resolving these relationships remains uncertain. Integrating cp genome data with nuclear DNA information offers a more comprehensive view but often requires separate datasets. In response, we employed the same raw read sequencing data to construct cp genome-based trees and nuclear DNA phylogenetic trees using Read2Tree, a cost-efficient method for extracting conserved nuclear gene sequences from raw read data, focusing on the Aurantioideae subfamily, which includes *Citrus* and its relatives. The resulting nuclear DNA trees were consistent with existing nuclear evolutionary relationships derived from high-throughput sequencing, but diverged from cp genome-based trees. To elucidate the underlying complex evolutionary processes causing these discordances, we implemented an integrative workflow that utilized multiple alignments of each gene generated by Read2Tree, in conjunction with other phylogenomic methods. Our analysis revealed that incomplete lineage sorting predominantly drives these discordances, while introgression and ancient introgression also contribute to topological discrepancies within certain clades. This study underscores the cost-effectiveness of using the same raw sequencing data for both cp and nuclear DNA analyses in understanding plant evolutionary relationships.

## Introduction

Phylogenetic analysis plays a crucial role in comprehending organismal evolution and taxonomy. Chloroplast (cp) genome sequences are widely utilized in plant phylogenetics due to their characteristics: typical maternal inheritance, stable gene content, low recombination, and relatively small genome size^1,2,3,4,5,6^. Typically spanning 120 to 170 kbp, the cp genome can often be assembled from as little as 1 Gbp of raw sequencing data^7,8^. However, sole reliance on cp genome sequences in plant phylogenetics has its limitations. Because cpDNA represents a single, uniparentally inherited genetic lineage, it evolves independently of the nuclear genome. This independence makes cpDNA particularly susceptible to genetic drift and lineage-specific events, such as chloroplast capture, where cpDNA is transferred between lineages through hybridization^9,10^. These events do not similarly affect the biparentally inherited nuclear genome, leading to discrepancies between phylogenetic signals from cpDNA and nuclear DNA, known as cytonuclear discordance^11,12,13^. Given these limitations, integrating cpDNA data with nuclear DNA data can provide a more comprehensive approach to phylogenetic reconstruction.

Nuclear DNA information, characterized by biparental inheritance and relatively high substitution rates, provides a more comprehensive genetic perspective than cpDNA^14,15^. However, the high cost of whole nuclear genome sequencing has often constrained its accessibility^16,17,18^. Recent technological advancements have mitigated these limitations by introducing cost-effective methods for extracting extensive nuclear loci^19,20^. Techniques such as genome skimming, target capture, and RNA sequencing (transcriptomics) are now central to phylogenetic studies, offering efficient avenues for generating nuclear data^20^. Furthermore, computational tools like Read2Tree, PhyloHerb, SECAPR, and HybPiper have made it easier to work with raw sequencing reads by tackling key challenges, including identifying genes, aligning sequences, and handling incomplete or low-quality data^21,22,23,24^.

Among these computational tools, the recently developed Read2Tree tool may represent a significant advancement in nuclear phylogenetics. It enables the precise extraction of numerous nuclear genes directly from raw sequencing reads through a reference-guided approach. Unlike other tools such as PhyloHerb, SECAPR, and HybPiper, which often require species-specific probe sets, custom-designed target gene databases, or separate workflows for different genomic compartments, Read2Tree integrates data extraction, alignment, and tree construction into a cohesive pipeline^21,22,23,24^. By leveraging predefined reference datasets, it eliminates the need for extensive validation of custom datasets. Furthermore, although it has not yet been attempted, the same sequencing reads may be utilized for both cp genome analysis and nuclear phylogenetic inference, enhancing overall efficiency.

Read2Tree produces two primary outputs: concatenated alignment data for tree inference and multiple alignments for each gene across the analyzed species. The concatenated alignment data is a primary focus of Read2Tree’s publication and has been used to reconstruct the phylogeny of yeast and various coronaviruses^21^. However, only one study has applied Read2Tree to plant phylogenetics^25^, and it did not validate Read2Tree’s results as comprehensively as the original publication^21^. Therefore, further research is needed to confirm Read2Tree’s effectiveness in plant phylogenetics and to compare its results with other nuclear DNA-based approaches.

The second output type of Read2Tree, multiple alignments for each gene, may present a promising approach to addressing the complexity of nuclear DNA evolution. In plants, interspecies hybridization occurs frequently, contributing to this complexity and serving as one of the key reasons why discordances between the species tree and gene trees are often observed^26,27^. These discordances are commonly attributed to genetic differentiation processes such as incomplete lineage sorting (ILS), introgression, and ancient introgression^28,29,30,31,32,33^. Methods leveraging multiple alignments for each gene are available to analyze these discordances and assess the effects of ILS, introgression, and ancient introgression. However, no studies have yet attempted to apply such analyses using the second output type of Read2Tree.

This study aims to elucidate and harness the potential of assembling nuclear datasets using Read2Tree in advancing phylogenetic reconstructions and discordance analysis, focusing on the phylogenetic relationships within Aurantioideae, a subfamily of Rutaceae that encompasses *Citrus* and its relatives. Previous research successfully used RAD-Seq to generate nuclear DNA data to resolve relationships in this subfamily^34^. While RAD-Seq focuses on nuclear DNA near specific restriction sites, Read2Tree assembles sequences from a broader set of conserved nuclear genes. By comparing the nuclear gene trees generated by Read2Tree with RAD-Seq results, we can more effectively evaluate the phylogenetic relationships. Second, the genus *Citrus*, along with other members of Aurantioideae, holds global importance due to its nutritional value, high vitamin C content, and widespread culinary use. Understanding the phylogenetic relationships within this group has implications not only for evolutionary biology but also for agriculture and conservation efforts. Many Aurantioideae species are valued for their edible and medicinal properties across various regions, making accurate phylogenetic reconstructions crucial for biodiversity preservation and resource management^35,36^.

In this study, we employed Read2Tree to assemble a large nuclear sequence dataset, which we used to investigate phylogenetic relationships among approximately 40 plant species within the subfamily Aurantioideae. We integrated nuclear phylogenies with whole cp genome analyses to explore evolutionary relationships more precisely and to investigate the potential causes of cytonuclear discordance. By comparing the nuclear phylogenetic trees generated by Read2Tree with those obtained from previous RAD-Seq analyses, we aimed to validate the robustness and utility of Read2Tree in plant phylogenetics. Additionally, we developed a comprehensive workflow that integrates Read2Tree with other phylogenomic methods to analyze both species-gene tree discordance and cytonuclear discordance. This workflow enabled us to assess whether the observed discordances were due to factors such as ILS, introgression, or ancient introgression followed by ILS. By identifying the sources of these discordances, we aimed to gain insights into the complex evolutionary history of Aurantioideae and the potential mechanisms driving species-gene tree discordance and cytonuclear discordance in plant phylogenetics.

## Results

### The phylogenetic tree constructed using Read2Tree demonstrates strong concordance with the tree derived from RAD-Seq data

The samples utilized in this investigation are detailed in Supplementary Table S1. Initially, Read2Tree facilitated the retrieval of sequences from various conserved nuclear genes. To establish the root of the phylogenetic tree within the subfamily Aurantioideae, a Maximum Likelihood (ML) tree was generated for 43 species of the Rutaceae family (Supplementary Fig. S1). Subsequently, we focused on 39 Aurantioideae species to generate an ML phylogenetic tree using the conserved nuclear gene sequences (Fig. 1), with rooting determined from the Rutaceae analysis. The ML phylogenetic trees largely aligned with findings from a prior RAD-Seq based study^34^, with noteworthy exceptions observed for *Micromelum minutum* and *Murraya koenigii*. Their phylogenetic positions, previously uncertain in the prior study^34^, remained unresolved in our current analysis, as evident from the comparison between Fig. 1 and Supplementary Fig. S1.

**Figure 1.**
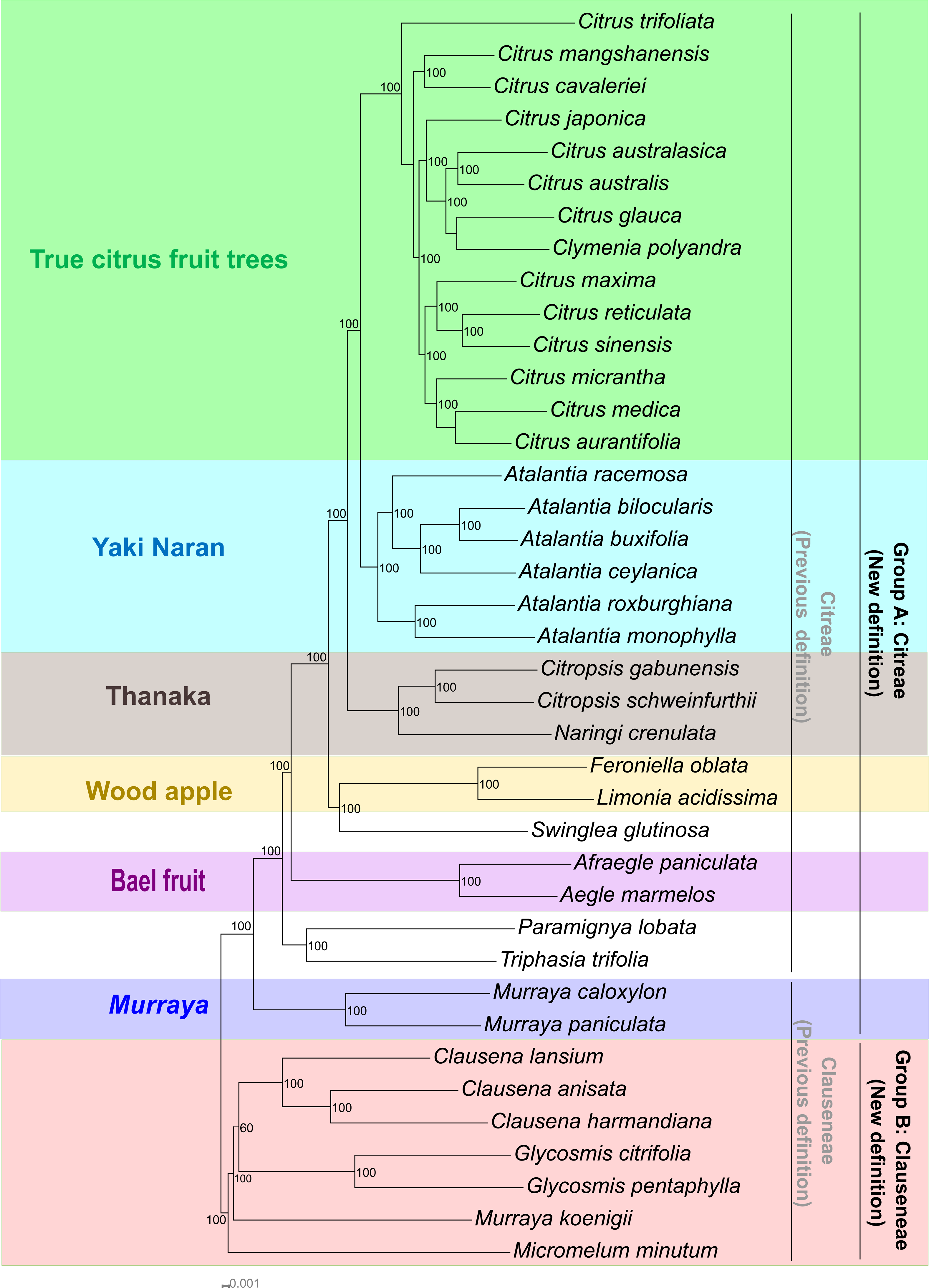
ML phylogenetic tree based on conserved nuclear gene sequences of 39 species in the Aurantioideae subfamily. The alignment used for phylogenetic tree construction included a total of 6,674,283 nucleotide positions with 605,597 parsimony-informative sites, as calculated using custom Python scripts (Supplementary Dataset 1). For the analysis, the optimal evolutionary model, GTRGAMMAIX, was employed in RAxML. The root position corresponded to that identified in the phylogenetic tree of the Rutaceae family (Supplementary Fig. S1). Bootstrap values (percentages over 100 replicates) are displayed at the nodes, with the scale bar indicating genetic divergence in substitutions per site. Bootstrap values below 50% are omitted. The subgroups identified in this study are represented by different colors. The alignment data in FASTA format and tree data in Newick format are available in Supplementary Dataset 2.

### Phylogenetic analysis employed whole cp genome sequences revealing disparities when compared with nuclear gene-based trees

Initially, a cp phylogenetic tree encompassing 59 members of the Rutaceae family was constructed, to determine the root position for subsequent analyses (Supplementary Fig. S2). Subsequently, cp phylogenetic trees were developed based on DNA sequences for the same 39 members of the Aurantioideae subfamily used in Read2Tree, utilizing ML and Bayesian Inference (BI) methods (Supplementary Fig. S3 and Supplementary Fig. S4, respectively). The inclusion of identical taxa in both the whole cp genome and nuclear analyses was a deliberate approach to minimize discrepancies arising from taxon sampling. The topologies of these cp phylogenetic trees remained consistent across the two analyses.

Comparison of the topologies of the two phylogenetic trees, one from whole cp genome sequences and the other from the concatenated nuclear gene tree generated by Read2Tree (Fig. 2A), revealed two primary groups within the Aurantioideae subfamily, referred to as group A and group B. However, this comparison detected cytonuclear discordances within each group (Fig. 2A). Specifically, inconsistencies were observed among the genera *Citrus*, *Atalantia*, and *Clausena*. Additionally, significant differences were found in intergeneric relationships. For instance, the cpDNA-based analysis positioned the *Limonia*/*Feroniella* subgroup as one of the closest outgroups to *Citrus*, contrary to the nuclear DNA analysis, which identified this subgroup as the nearest outgroup to a clade comprising *Citrus, Atalantia, Citropsis*, and *Naringi*. Furthermore, the cpDNA analysis grouped *Swinglea* with *Citropsis* and *Naringi*, while nuclear DNA analysis aligned *Swinglea* with the *Limonia*/*Feroniella* subgroup. Discrepancies between the two analyses were also evident in the phylogenetic structures formed by *Paramignya*, *Triphasia*, and *Aegle*/*Afraegle*. The phylogenetic positions of *Micromelum minutum* and *Murraya koenigii* remained ambiguous.

**Figure 2.**
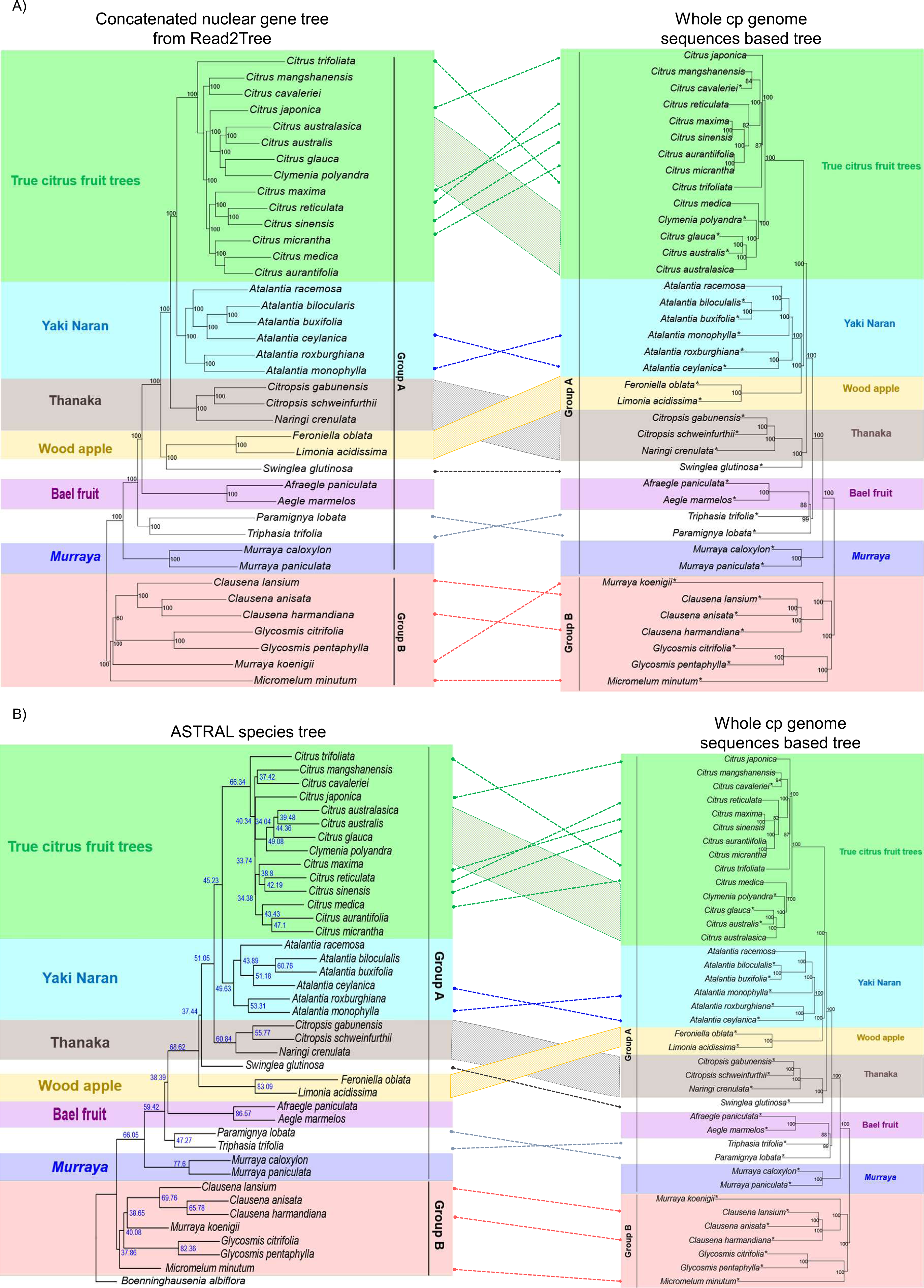
Discordance between nuclear gene-derived trees and cp tree (A) Comparison of the concatenated nuclear gene tree and the cp tree. Left panel: ML phylogenetic tree generated using Read2Tree, based on conserved nuclear gene sequences from 39 species within the Aurantioideae subfamily. Bootstrap support values (percentages from 100 replicates) are displayed at each node, with values below 50% omitted. Subgroups identified in this study are represented by distinct colors. (B) Comparison of the ASTRAL species tree and the cp tree. Left panel: Species tree inferred using ASTRAL-III, with local posterior probability (LPP) values displayed as percentages for each corresponding node. Subgroups identified in this study are represented by different colors. Tree data in Newick format are available in Supplementary Dataset 4. Right panel in both A and B: ML phylogenetic tree constructed using the whole cp genome sequences of 39 species in the Aurantioideae subfamily. Bootstrap support values (percentages from 1,000 replicates) are displayed at the nodes, with values below 50% omitted. Asterisks mark sequences that were assembled de novo in this study. Subgroups identified are represented by distinct colors. Dotted lines and shapes highlight conflicting relationships between the nuclear and cp phylogenies, revealing areas of topological incongruence between the two trees. Alignment data (FASTA format) and tree data (Newick format) are available in Supplementary Dataset 3.

### Nuclear gene assembly from Read2Tree and insights into phylogenetic discordances

The detection of these discordances strongly indicates the presence of broader phylogenetic incongruences. By integrating Read2Tree with other phylogenomic methods, these discordances were determined. Orthologous groups (OGs), consisting of genes or DNA sequences in different species that diverged from a common ancestral gene through speciation, are critical for studying evolutionary relationships^37^. In this analysis, Read2Tree generated multiple sequence alignments for each OG, which were used for constructing each gene tree. For this analysis, in addition to the 39 members of the Aurantioideae subfamily used in the aforementioned analysis, *Boenninghausenia albiflora* was employed as the sole outgroup to maximize the number of available multiple sequence alignments for each OG. Specifically, alignments from 579 OGs with a ratio of N characters to total nucleotides below 10% were identified through Read2Tree analyses (Supplementary Table S2). The number of genes successfully recovered varied between samples, ranging from 511 genes for *Boenninghausenia albiflora* to 576 genes for *Citrus trifoliata*, with missing data consistently under 20% for each sample (Supplementary Table S3). Furthermore, the recovered data exceeded 80% in all samples, and we observed a positive correlation between the number of paired reads and the percentage of recovered data (Supplementary Table S3). However, this relationship was weak, likely influenced by additional factors such as variations in genome size and structural complexity (Supplementary Table S3). In particular, the relationship with genome size is an intriguing issue. However, due to discrepancies observed between genome size estimates based on k-mers and the known genome size, we decided not to perform genome size estimation.

The 579 gene trees were used to infer the species tree through the ASTRAL method, as visualized in Fig. 2B (left panel) and Fig. 3A (right panel). Additionally, a concatenated nuclear gene tree based on this subset of 579 genes was generated (Supplementary Fig. S5, right panel). The resulting species tree yielded a normalized quartet score (QT) of 0.7, indicating that 70% of the quartets in the gene trees were concordant with the species tree. A comparison of the two trees revealed discrepancies only within the genus *Citrus* and *Micromelum minutum* (Supplementary Fig. S5 B). Within *Citrus*, the observed discrepancies were associated with low bootstrap support (<50), suggesting that species positioning within this genus is unreliable in the concatenated nuclear gene tree based on the 579-gene subset. The placement of *Micromelum minutum* also remains uncertain. Since *Boenninghausenia albiflora* was used as the sole outgroup, the accuracy of the root position is uncertain. However, both trees suggest the presence of two groups, Group A and Group B. This observation agrees with the groupings identified in the cp phylogeny and the concatenated nuclear gene phylogeny generated using Read2Tree, as shown in Fig. 2B, Fig. 3A, and Supplementary Fig. S5.

**Figure 3.**
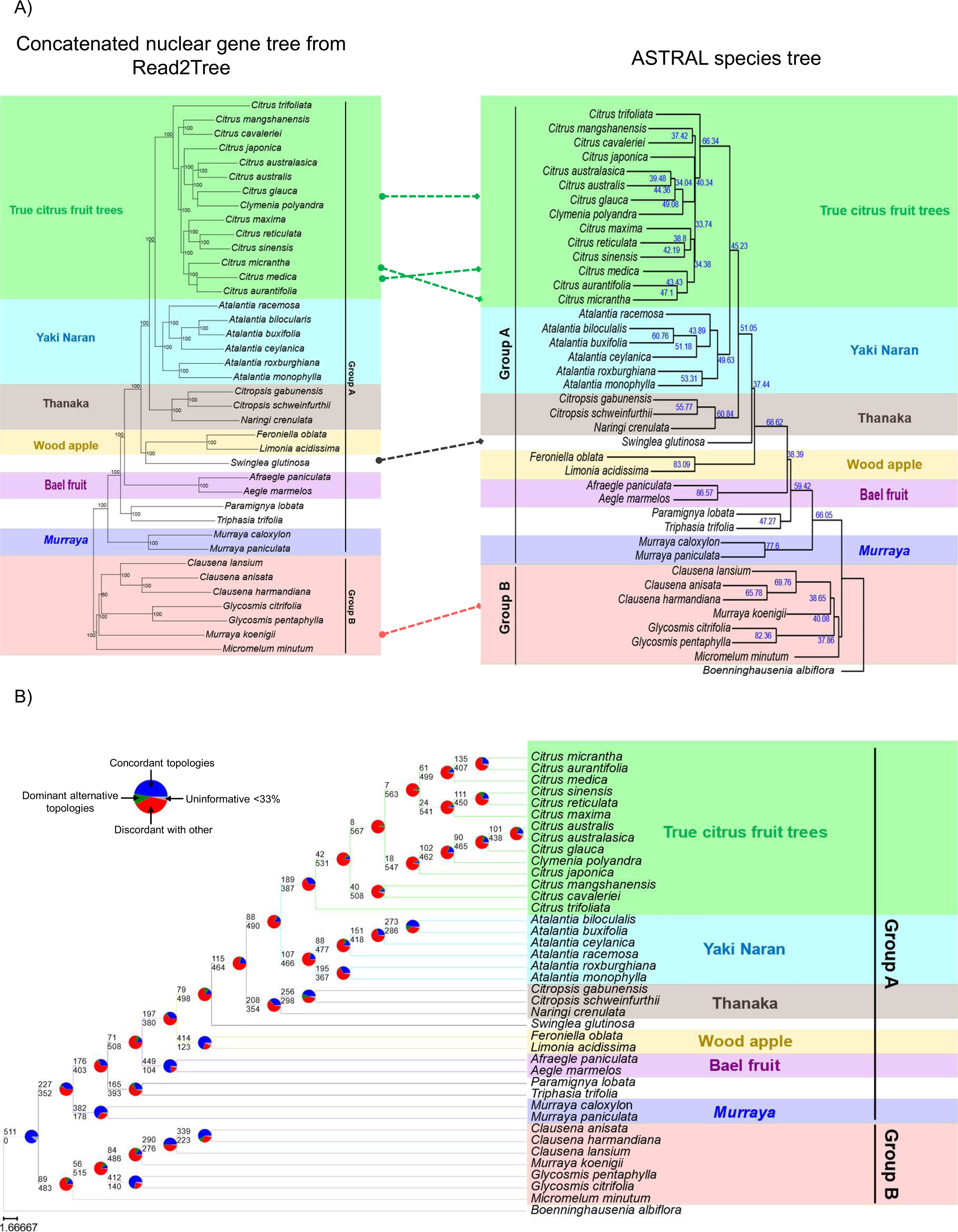
Species tree and concatenated nuclear gene tree (A) Comparison of the concatenated nuclear gene tree and ASTRAL species tree. Left panel: ML phylogenetic tree generated using Read2Tree, based on conserved nuclear gene sequences from 39 species within the Aurantioideae subfamily. Bootstrap support values (percentages from 100 replicates) are displayed at each node, with values below 50% omitted. Subgroups identified in this study are highlighted in distinct colors. Right panel: Species tree inferred using ASTRAL-III, with local posterior probability (LPP) values displayed as percentages for each corresponding node. Subgroups identified in this study are denoted by distinct colors. Dotted lines indicate areas of incongruence between the two trees. (B) Gene tree discordance displayed on the ASTRAL-inferred species tree. Pie charts at each node illustrate the proportion of gene trees falling into one of four categories: concordant with the species tree (blue), discordant but representing the most common alternative topology (green), discordant with other alternative topologies (red), and uninformative due to low branch support (gray, <33% bootstrap support at the node). The number above each branch indicates the count of gene trees concordant with the species tree topology at that node (blue portion of the pie). The number below each branch represents the sum of informative discordant topologies at that node (green + red portions of the pie). Different colors denote the subgroups identified in this study.

When comparing the species tree to the concatenated nuclear gene tree constructed using Read2Tree (Fig. 3A), notable discrepancies are observed within the *Citrus* genus and in the intergeneric relationships involving *Swinglea* and *Murraya koenigii* (Fig. 3A). In the species tree, *Swinglea* is placed as an outgroup to *Citrus*, *Atalantia*, and *Citropsis/Naringi*, whereas in the concatenated tree, *Swinglea* clusters with *Limonia* and *Feroniella*. Additionally, *Murraya koenigii* acts as a near outgroup exclusively to *Clausena* species in the species tree, while in the concatenated tree, it serves as an outgroup to both *Clausena* and *Glycosmis* species. This ambiguous positioning of *Murraya koenigii* mirrors the unresolved status observed in previous RAD-Seq studies as well as in our Read2Tree and cp genome trees (Fig. 2A).

PhyParts conflict analysis was performed to investigate the complex discordances between gene trees and the species tree, with *Boenninghausenia albiflora* used as the outgroup (Fig. 3B). The analysis revealed that 511 out of 579 gene trees were concordant with the topology supporting the presence of two primary groups, groups A and B, within the subfamily Aurantioideae. However, significant conflict was observed among informative gene trees at the node encompassing *Citrus*, *Atalantia, Citropsis/Naringi*, *Swinglea*, *Paramignya*/*Triphasia*, *Murraya koenigii*, and *Micromelum minutum*.

As mentioned above, cytonuclear discordance was observed between the concatenated nuclear gene tree constructed using Read2Tree and the cp tree (Fig. 2A). Additionally, significant cytonuclear discordances were evident between the species tree and the cp tree, particularly within the *Citrus*, *Atalantia*, and *Clausena* genera, as well as in intergeneric relationships (Fig. 2B). The species tree positions the *Limonia/Feroniella* subgroup as an outgroup to a clade containing *Citrus*, *Atalantia, Citropsis/Naringi*, and *Swinglea*, while the cp tree places this subgroup as the closest outgroup to *Citrus* and *Atalantia*. Furthermore, in the species tree, *Swinglea* is positioned as an outgroup to *Citrus, Atalantia, Citropsis*, and *Naringi*, but the cp tree groups *Swinglea* with *Citropsis/Naringi*. Discrepancies between the species tree and the cp tree are also apparent in the phylogenetic structures of *Paramignya, Triphasia*, and *Aegle/Afraegle*. Additionally, in the ASTRAL species tree, *Micromelum minutum* is positioned as an outgroup to both *Clausena* and *Glycosmis*, whereas in the cp tree, it appears as a near outgroup exclusively to *Glycosmis*.

### Consistent subgrouping patterns unveiled by nuclear and cp trees

Both nuclear and cp analyses demonstrated similar grouping and subgrouping patterns, as depicted in Figures 2, 3, and Supplementary Figure S5. The phylogenies derived from both nuclear and cp data consistently revealed two primary groups, designated as group A and group B, within the subfamily Aurantioideae (Figs. 1, 2, 3, and Supplementary Fig. S5). Historically, Aurantioideae has been classified into the Citreae and Clauseneae tribes^38^ (Supplementary Table S4). However, our analysis led to a reassignment of species within these tribes. Specifically, *Murraya caloxylon* and *Murraya paniculata*, historically classified within the Clauseneae tribe, have been reassigned to the Citreae tribe, now defined as group A. The remaining species historically classified under the Clauseneae tribe have been consolidated into group B, which corresponds to the newly defined Clauseneae tribe in this study (Fig. 2 and Fig. 3).

Within group A, six distinct subgroups were identified. The “True citrus fruit trees” subgroup comprises *Citrus* species, consistent with traditional taxonomy^38^. The “Yaki Naran” subgroup encompasses all *Atalantia* species, inspired by Ayurvedic medicine in Sri Lanka, including *Atalantia ceylanica*. The “Thanaka” subgroup, named after the traditional cosmetic from Myanmar, comprises *Citropsis* and *Naringi* species. The “Wood apple” subgroup includes *Limonia* and *Feroniella* species, while the “Bael fruit” subgroup comprises *Aegle* and *Afraegle,* reflecting their historical names^38^. The final subgroup, the *Murraya* subgroup, consists of *Murraya caloxylon* and *Murraya paniculata*. However, three species *Swinglea glutinosa*, *Paramignya lobata*, and *Triphasia trifolia*, did not fit neatly into any defined subgroup.

### Signatures of ILS, introgression, and ancient introgression

Discordances were detected among three key groups: (1) True citrus fruit trees and its near subgroups, (2) the Bael fruit subgroup and its associated species, and (3) Group B (Fig. 4). These discordances likely arise from multiple sources, including errors and noise in data assembly and filtering, hidden paralogy, random noise from uninformative genes, and processes associated with genetic differentiation, such as ILS, introgression, and ancient introgression. Among these factors, the genetic differentiation processes, particularly ILS and introgression, have been most extensively studied. To investigate the impact of these processes on the observed discordance patterns, one representative species from each subgroup was selected for detailed analysis. Introgression was assessed using Patterson’s D-statistic (significance threshold: P < 0.05)^39^, while D_FOIL_ analysis was applied to detect ancient introgression events (significance threshold: P < 0.05)^40^. To further disentangle the contributions of ILS and introgression to the observed phylogenetic discordances, the Quantifying Introgression via Branch Lengths (QuIBL) method was employed^41^. This method evaluates triplet topologies from gene trees by comparing branch length distributions under two models: one accounting for ILS alone and the other incorporating both ILS and introgression. Model selection was guided by Bayesian Information Criterion (BIC) values, with ΔBIC used to differentiate between ILS and introgression. Specifically, a ΔBIC greater than 10 indicates ILS, a value lower than −10 suggests introgression, and values within this range imply that ILS and introgression cannot be clearly distinguished.

**Figure 4.**
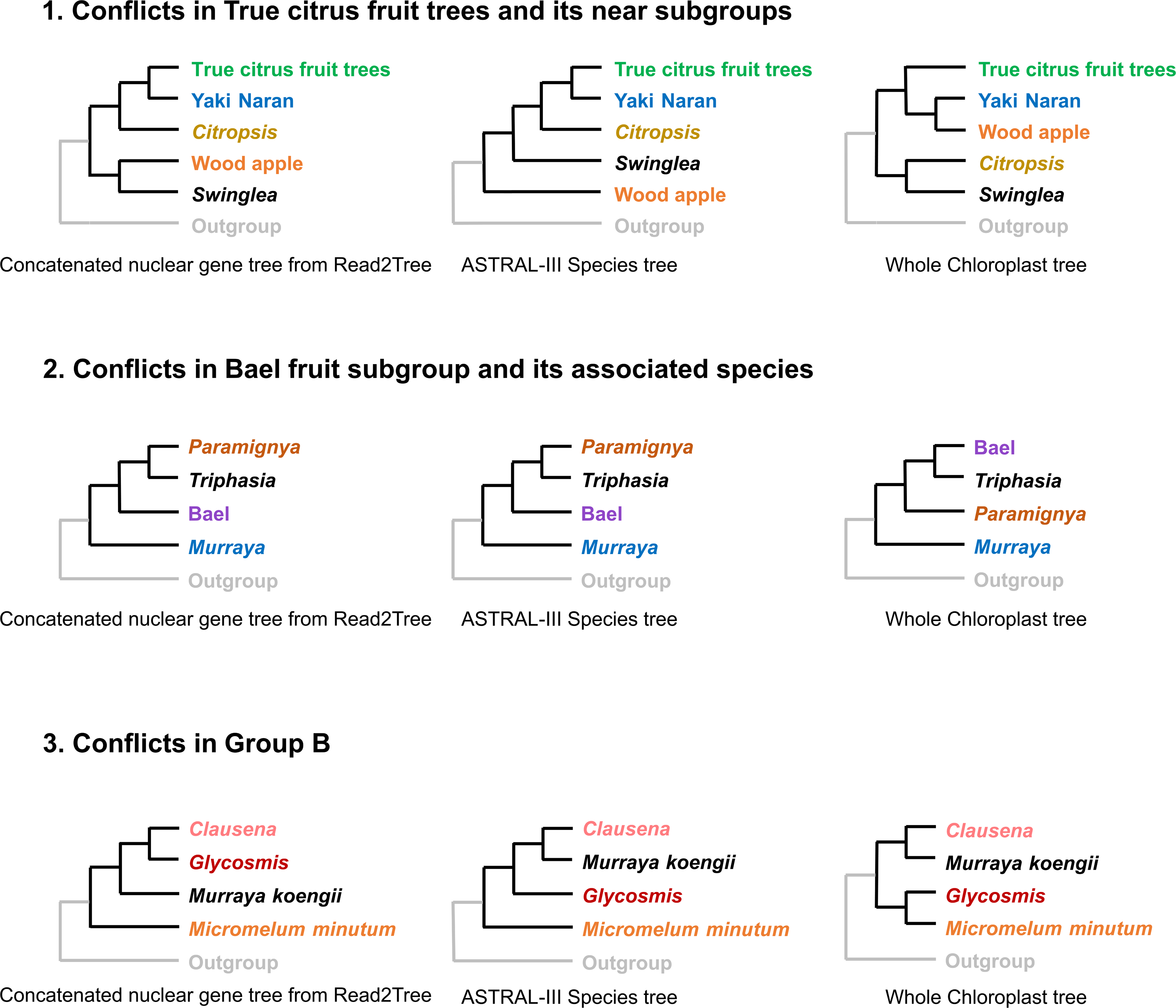
Discordance analysis among three subgroups: (1) True citrus fruit trees and its near subgroups, (2) Bael fruit subgroup and its associates, and (3) the Group B. One representative from each subgroup/genus is used.

In the True citrus fruit trees and its near subgroups, we analyzed 860 gene trees to investigate patterns of phylogenetic discordance. Patterson’s D-statistic identified significant introgression events between True citrus fruit trees and Yaki Naran, as well as between Yaki Naran and Wood apple (Supplementary Table S5). D_FOIL_ analysis further suggested that discordant topologies within this group were largely influenced by ancient introgression followed by ILS (Supplementary Table S6). To disentangle the roles of ILS and introgression, QuIBL analysis was performed on 30 triplets (Supplementary Table S7). Among these, 33% (10 of 30) showed evidence of ILS (ΔBIC > 10), 10% (3 of 30) supported introgression (ΔBIC < −10), and 26.67% (8 of 30) were inconclusive (ΔBIC between −10 and 10). Specifically, the discordant topology involving wood apple, *Swinglea glutinosa*, and *Citropsis gabunensis* (used as an outgroup) was attributed to introgression (ΔBIC < - 10). Furthermore, PhyloNet analysis provided strong support for introgression events within the True citrus fruit trees and its near subgroups (Fig. 5B and Supplementary Fig. S6). These results suggest that ancient introgression followed by ILS is the dominant factor contributing to discordances in True citrus fruit trees and its near subgroups.

**Figure 5.**
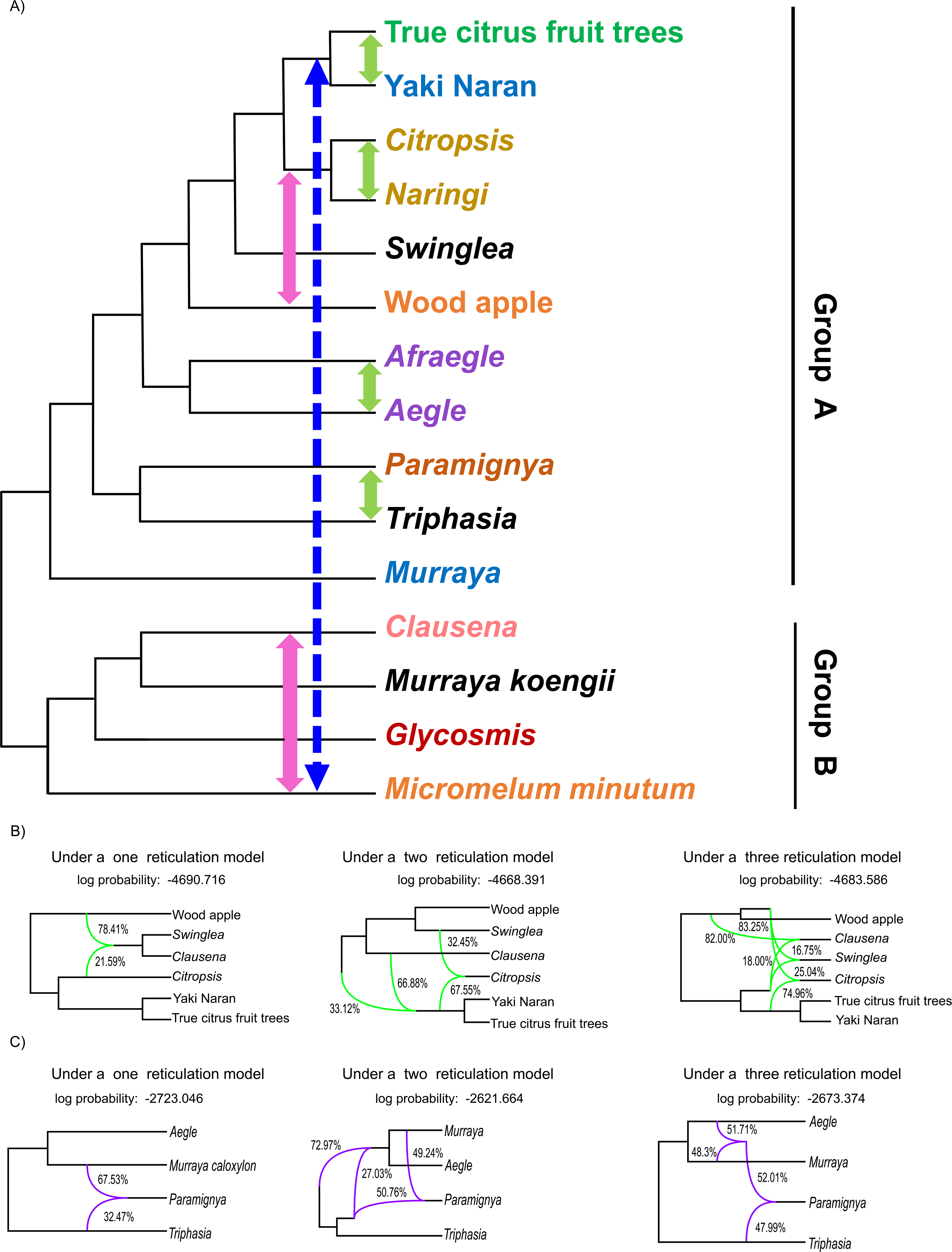
Summary of phylogenetic discordance sources (A) ASTRAL-inferred species tree depicting the relationships among members of the Aurantioideae subfamily. Arrows indicate various evolutionary events: the dashed blue arrow represents ILS; green arrows represent introgression; and pink arrows indicate ancient hybridization followed by ILS. Detailed results are provided in Supplementary Tables 5 to 14. One representative from each genus/subgroup is used. Phylogenetic networks for True citrus fruit trees and its near subgroups (B), as well as the Bael fruit subgroup and associated species (C), were inferred using PhyloNet with Maximum Pseudolikelihood under models allowing for one, two, and three reticulations. The Akaike Information Criterion (AIC) indicates that the three-reticulation model is the best fit for both groups. In Figure (B), green branches highlight lineages involved in reticulate evolutionary histories, with numerical values representing the inheritance probabilities for each reticulation event. In Figure (C), purple branches similarly indicate lineages associated with reticulation, along with their corresponding inheritance probabilities. The Nexus files for each reticulation model can be found in Supplementary Datasets 5 and 6. Supplementary Figs. S6 and S7 present all inferred phylogenetic networks generated by PhyloNet for True citrus fruit trees and its near subgroups and the Bael fruit subgroup with associated species, respectively.

The analysis of the Bael fruit subgroup and its associated species was done based on 1,964 gene trees. Patterson’s D-statistic and PhyloNet analyses identified introgression between *Paramignya lobata* and *Triphasia trifolia* as a significant contributor to the observed inconsistencies (Fig. 5C, Supplementary Fig. S7, and Supplementary Table S8). Additionally, QuIBL analysis revealed that ILS was responsible for the discordant topologies in 66.67% of cases. However, the triplet involving *Paramignya lobata*, *Triphasia trifolia*, and *Aegle marmelos* (used as an outgroup) produced ΔBIC values between −10 and 10, suggesting that the discordance in this specific case could be attributed to either ILS or introgression (Fig. 5A and Supplementary Table S9). These results indicate that while introgression has contributed to specific instances of phylogenetic discordance, particularly between *Paramignya lobata* and *Triphasia trifolia*, ILS accounts for the majority of discordant topologies within the Bael fruit subgroup and its associated species.

Within Group B, analyses of 1,135 gene trees using Patterson’s D-statistic and QuIBL indicated that ILS accounted for a significant proportion of the discordant topologies. QuIBL revealed evidence of ILS in 58.33% of triplets (7 out of 12, ΔBIC > 10), while 16.67% (2 out of 12, ΔBIC < - 10) supported introgression, and 25% (3 out of 12) were ambiguous (ΔBIC between −10 and 10). In contrast, D_FOIL_ analysis identified signatures of ancient introgression involving *Glycosmis citrifolia* and *Micromelum minutum* with other taxa (Fig. 5A). These results suggest that discordance within Group B is largely explained by ancient introgression followed by ILS (Supplementary Tables S10, S11, and S12).

The source of cytonuclear discordance was inferred using D-statistics and QuIBL results. The results suggested that ILS is a significant driver of the observed cytonuclear discordance within True citrus fruit trees and its near subgroups, as well as the Bael fruit subgroup and its associates (Supplementary Tables S5, S7, S8, and S9). Additionally, both D_FOIL_ and QuIBL analyses revealed that the discordant topologies involving *Micromelum minutum* and *Murraya koenigii* are predominantly due to ancient introgression, followed by ILS (Supplementary Tables S10, S11, and S12).

### Introgression events between Asian and African plants

The generated phylogenetic trees revealed two instances of close relationships between Asian and African plants in the Thanaka and Bael fruit subgroups (Fig. 2, Fig. 3 and Supplementary Fig. S5). Within the Thanaka subgroup, *Citropsis* is native to Africa, while *Naringi* originates from Asia. Similarly, within the Bael fruit subgroup, *Aegle* is indigenous to Asia, and *Afraegle* is native to Africa. To investigate the discordance in topologies involving *Citropsis* and *Naringi*, we analyzed a total of 1,084 gene trees. Similarly, we evaluated 1,080 gene trees to explore the discordance in topologies involving *Afraegle* and *Aegle*. Our Patterson’s D-statistic and QuIBL analyses revealed compelling evidence of introgression events between these Asian and African plant lineages (Fig. 5A, Supplementary Tables S13 and S14). Specifically, the introgression event within the Bael fruit subgroup occurred over 1 to 2 coalescent time scales, while the introgression event in the Thanaka subgroup occurred over 0.5 to 2 coalescent time scales (Supplementary Table S14).

## Discussion

In this study, we leveraged an accessible sequence dataset to assemble and analyze a large number of nuclear gene sequences alongside whole cp genome sequences for phylogenetic reconstructions. Rather than relying on a single data type, the integration of both nuclear and cp genome sequences offers a more comprehensive and informative perspective in plant phylogenetics^12,13,42,43,44^. Unlike previous studies that used separate datasets for nuclear and cp genome sequences^42,43,44^, our approach is more cost-efficient as we used the same short-read sequence data for both analyses.

Our analysis revealed discrepancies between phylogenetic trees derived from nuclear gene sequences and those based on whole cp genome sequences. Despite these differences, all trees consistently identified two primary groups with similar subgrouping patterns. These results underscore the need for reclassification within the Aurantioideae subfamily to better align with the evolutionary relationships revealed by both nuclear and cp genome data. In response, we propose a reclassification that refines the taxonomy of Aurantioideae, providing a more accurate reflection of the subfamily’s genetic and evolutionary relationships.

The nuclear sequences assembled using Read2Tree are useful not only for core phylogenetic reconstruction, but they also can be used to address phylogenetic discordance. It enables the direct acquisition of a single multiple sequence alignment for each OG, which is crucial for constructing individual gene trees. Data from these gene alignments can be used with other phylogenomic tools to identify evidence of processes such as ILS, introgression, and ancient introgression that contribute to discordances. These evolutionary processes pose challenges to phylogenomics by complicating the inference of accurate evolutionary relationships^11,30,31,32,33^. ILS arises when ancestral polymorphisms persist through successive speciation events, especially during rapid radiation with short intervals between these events^30,31,32,33,45,46^. This persistence can lead to discordant gene tree topologies. Introgression, which involves the transfer of genetic material between species through hybridization, can blur species boundaries and distort gene trees^28,31,32,46,47^. Ancient introgression specifically refers to gene flow that took place in the distant past, often during the early branching events of a phylogenetic tree, involving species that have diverged for considerable periods^28,29,30,31^.

In addition to genetic differentiation processes, errors in gene tree inference including stochastic errors, systematic errors, and issues related to paralogy can substantially contribute to observed discordances in phylogenetic analyses^45,48^. To address these challenges and improve the reliability of our findings, we implemented multiple strategies aimed at mitigating these errors. To reduce the impact of stochastic and systematic errors, we utilized 100 bootstrap replicates to bolster the robustness of gene tree topologies and conducted thorough model selection for each gene tree using ModelTest-NG, optimizing the application of evolutionary models. Additionally, we applied the coalescent-based ASTRAL method to reconcile individual gene trees with the species tree, enhancing the accuracy of the overall phylogenetic framework. We also took measures to manage issues related to paralogy, which can introduce misleading phylogenetic signals by conflating homologous genes with gene duplications. The Read2Tree method, specifically designed to minimize paralogy^21^, was effectively employed in our study, reducing its impact on the constructed gene trees. Furthermore, because the analyzed species are not estimated to be polyploid, the risk of paralogous gene duplications confounding the phylogenetic signal was minimized. Overall, our analysis strongly suggests that ILS drives most of the discordances we observed. This is consistent with the rapid diversification within the Aurantioideae subfamily, which has obscured true phylogenetic relationships and complicated the reconstruction of its evolutionary history.

Our discordance analysis concentrated on three key groups: True citrus fruit trees and its near subgroups, the Bael fruit subgroup and its associated species, and group B. Some of the discordant topologies in these areas were consistent with introgression and ancient introgression, followed by ILS. These introgressive events may have facilitated gene flow between distinct lineages, resulting in discrepancies between gene trees and the species tree. Additionally, ILS preserves ancestral genetic polymorphisms, further complicating the reconstruction of accurate phylogenetic histories^30,31,42,46,49,50^. In the context of True citrus fruit trees and its near subgroups, our results align with previous studies that reported hybridization between *Citrus* and intergeneric species, emphasizing the substantial impact of introgression on shaping phylogenetic relationships within this group^51,52^. Prior research also highlights how hybridization frequently leads to incongruence among gene trees, underscoring its importance in accurately inferring the phylogeny of *Citrus* species^53^.

Topological conflicts between nuclear and organellar phylogenies, known as cytonuclear discordance, are common in phylogenetic studies^12,13,42,43,44,54,55^. Our analyses suggest that ILS is more likely to contribute to cytonuclear discordance than introgression. Rapid speciation events within Aurantioideae may have occurred with insufficient time for gene lineages to coalesce, resulting in discordant gene trees between nuclear and organellar genomes^56^. Additionally, large effective population sizes can also extend the persistence of multiple gene lineages, increasing the likelihood of discordant phylogenies^56,57^. While historical hybridization and introgression introduce additional genetic material and create a mosaic of gene trees^58,59,60^, our findings suggest that ILS plays a primary role in the observed cytonuclear discordance, underscoring the complex evolutionary history of the Aurantioideae subfamily.

The concatenated nuclear gene tree derived from Read2Tree data shows strong congruence with the RAD-seq phylogenetic tree, suggesting consistent evolutionary relationships (Fig. 1). In contrast, the ASTRAL species tree and the concatenated tree constructed from a dataset of 579 genes present different perspectives (Supplementary Fig. S5). The former analysis uses relatively larger datasets but contains many missing values, whereas the latter employs relatively smaller datasets with fewer missing values. These observations highlight the distinct strengths and limitations of each approach, making it challenging to determine the most accurate or optimal phylogenetic method.

In conclusion, our study highlights the effectiveness of the Read2Tree method in plant phylogenetics, showcasing its utility not only in reconstructing phylogenies but also in addressing phylogenetic discordance. By integrating Read2Tree with other phylogenomic approaches, we were able to identify key genetic differentiation processes, including ILS, introgression, and ancient introgression, which contribute to phylogenetic conflicts. This sheds light on the method’s robustness and versatility, advancing our understanding of complex evolutionary relationships and offering a valuable tool for future research. Additionally, our findings emphasize the critical importance of complete data submission for ensuring the transparency, reproducibility, and robustness of phylogenetic analyses. While many studies have submitted only assembled whole cp genome sequences to public databases, the inclusion of the raw sequence data used in the assembly process is essential, especially for methods like Read2Tree that rely on these data. Ensuring the availability of raw data allows for more comprehensive analyses, facilitating validation, replication, and deeper insights into evolutionary relationships.

## Methods

### DNA sequence data collection and processing

The DNA used in this study was either sourced from the same batch used in a previous experiment^34^ or purified using the method described therein. DNA quality was assessed by electrophoresis on a 1% agarose gel, and DNA concentration was measured using a DNA BR Assay Kit (Thermo Fisher Scientific, USA). Sequencing libraries were prepared from total DNA samples by Novogene (Singapore), using the NEBNext Ultra DNA Library Prep Kit for Illumina (NEB, USA). These libraries were then sequenced to produce 150-bp paired-end reads using a NovaSeq 6000 instrument (Illumina, USA) by Novogene. Additional DNA sequence data were acquired from public databases, with accession numbers detailed in Supplementary Table S1. For preprocessing all DNA sequence data, FastP (version 0.23.2) was employed with its default settings^61^.

### Concatenated phylogenetic analysis based on nuclear gene sequences from Read2Tree

For the concatenated phylogenetic analysis, we used Read2Tree (version 0.1.5)^21^. Marker genes for *Citrus clementina*, *Eucalyptus grandis*, all five species of the Malvaceae, and all seven species of the Brassicaceae were obtained from the OMA browser (https://omabrowser.org/oma/export_markers). The OMA browser parameters were set with a minimum fraction of covered species at 0.8 and a maximum number of markers at –1. In total, 16,258 OGs were identified among the selected 14 species in the OMA browser. Read2Tree was utilized to generate alignment files using these marker genes. To ensure the specificity of the phylogenetic trees, the 14 species obtained from the OMA browser were excluded from the resulting alignment files. Aligned data were trimmed using trimAl (version 1.4. rev15) with the − automated1 option^62^. ModelTest-NG (version 0.1.7) was used to identify the optimal evolutionary model^63^. Subsequently, Maximum Likelihood (ML) trees were generated using RAxML (version 8.2.12) with 100 bootstrap replicates^64^. The visualization of the resulting trees was achieved using Dendroscope (version 3.8.8)^65^.

### *De novo* assembly and annotation of cp genomes

The *de novo* assembly of cp genomes was performed using GetOrganelle (version 1.7.5)^66^. The parameters for this process were set to − R 15, − k 21,45,65,85,105, and − F embplant_pt. To validate the assembly, we utilized Bowtie2 (version 2.4.4) for mapping^67^. The sorting of the mapped data and the generation of bam files were conducted using Samtools (version 1.14)^68^. Assembly quality was confirmed, and heteroplasmic sites were identified using the Integrated Genome Viewer^69^. Despite detecting heteroplasmic sites, the assemblies obtained from GetOrganelle were used for subsequent analysis.

For annotation of the assembled cp genomes, we used GeSeq^70^. Within the GeSeq framework, protein-coding sequences (CDS) were predicted using Chloë (version 0.1.0)^71^, tRNAs were identified using tRNAscan-SE (version 2.0.7)^72^, and rRNAs were predicted using HMMER^73^. Thereafter, the annotation results were curated manually. Circular genome maps of the annotated cp genomes were generated using OGDRAW (version 1.3.1)^74^.

### Phylogenetic analysis based on cp genome sequences

For phylogenetic analysis of cp genome sequences, DNA sequence alignments were performed using HomBlocks (version 1.0) with its default settings^75^. The optimal evolutionary model for the alignments was identified using ModelTest-NG (version 0.1.7)^63^. Subsequently, ML trees were constructed using RAxML (version 8.2.12), employing 1000 bootstrap replicates^64^. The visualization of the resultant ML trees was accomplished using Dendroscope (version 3.8.8)^65^.

For the construction of Bayesian Inference (BI) trees, BEAST (version 1.10.4) was used, implementing the Markov Chain Monte Carlo (MCMC) method over a total of 10,000,000 generations^76^. Tree construction samples were collected every 10,000 generations, with the initial 10% of generations discarded as ‘burn-in’ to reduce potential bias from the early stages of the MCMC run. The completed BI trees were visualized using FigTree (version 1.4.4) (http://tree.bio.ed.ac.uk/software/figtree/).

### Species tree inference from Read2Tree data

To construct the species tree, alignments for 16,284 OGs across all species were utilized from the Read2Tree output file “06_align_merge_dna”. The OG alignments were processed using trimAl (version 1.4, revision 15)^62^ with the -automated1 option and eliminate species present in the OMA browser. The alignments were then assessed based on the ratio of missing data (N characters) to total nucleotide characters, with a threshold set at 10%. From this, 579 OGs were selected for further analysis (Supplementary Dataset 10). Model testing^63^ was performed for each OG, and the ML gene tree of each OG was constructed using RAxML (version 8.2.12) with 100 bootstrap replicates^64^. The species tree was subsequently inferred using ASTRAL-III (version 5.7.8)^77^. Additionally, a concatenated ML nuclear gene tree based on the subset of 579 genes was constructed. This process involved sequence concatenation with catfasta2phyml (version 1.2.1), model testing using ModelTest-NG (version 0.1.7)^63^, and tree construction using RAxML (version 8.2.12) with 100 bootstrap replicates^64^. Visualization of the resulting ML trees was performed with Dendroscope (version 3.8.8)^65^.

### Phylogenetic discordance analysis

PhyParts was utilized to evaluate the concordance and discordance between gene trees and the species tree^78^. Gene trees were processed by collapsing branches with bootstrap support below 33% and subsequently rerooted using *Boenninghausenia albiflora* as the outgroup. Similarly, the ASTRAL species tree was rerooted with *B. albiflora*, and this rerooted tree served as the reference for the conflict analysis. The PhyParts output was visualized on the ASTRAL topology using the phypartspiecharts.py script available at github.com/mossmatters/phyloscripts. The analysis provided metrics on the number of gene trees that were concordant or discordant with the species tree topology at each branch, as well as those that were uninformative (i.e., gene trees with support values below 33% at the given branch).

The discordance analysis was focused on three specific issues: (1) *Citrus* and its near subgroups using 860 gene trees (Supplementary Dataset 12), (2) Bael fruit subgroup and its associated species using 1,964 gene trees (Supplementary Dataset 13), and (3) Group B using 1,135 gene trees (Supplementary Dataset 14), with one representative from each subgroup included. Gene trees for these analyses were generated using similar criteria and procedures as those applied in species tree estimation. Additionally, 1,084 gene trees were generated to assess discordance in topologies involving *Citropsis* and *Naringi* (Supplementary Dataset 15), while 1,080 gene trees were used to examine discordance involving *Afraegle* and *Aegle* (Supplementary Dataset 16). Patterson’s D- statistic analysis was performed using D-suite (version 0.4)^39^ with the Dtrios option to identify introgression events (P < 0.05). Additionally, D_FOIL_ statistics^40^ were employed to detect ancient introgression events, with significance determined by P < 0.05. QuIBL was employed to distinguish between ILS and introgression as causes of the observed discordances^41^. Model selection was guided by BIC values, with ΔBIC values interpreted as follows: ΔBIC > 10 indicates ILS only, ΔBIC < −10 suggests introgression, and −10 < ΔBIC < 10 indicates either ILS or introgression.

To further investigate reticulation events, Maximum Pseudolikelihood (MPL) analysis in PhyloNet (version 3.8.2)^79^ was conducted. We performed 10 runs per search, each returning five optimal networks. These networks were then optimized for branch lengths and inheritance probabilities under full likelihood, using default settings and the -po option. The best-fitting network was selected based on the Akaike Information Criterion (AIC)^80,81^, with the number of parameters (k) being the sum of branch lengths and reticulations, and L representing the likelihood value.

### Use of generative AI

ChatGPT (https://chat.openai.com/) was used to rewrite the manuscript into more appropriate English and proofread the English text. Subsequently, the authors verified the correctness of the revisions.

## Supporting information

Legends for Supplementary Figures

## Acknowledgements

In the initial phase of our study’s data analysis conducted in Sri Lanka; we extend our profound gratitude to Dr. Dhammika Elkaduwe from the Faculty of Engineering at the University of Peradeniya for his exceptional support in granting access to high-performance computing facilities. We would like to thank Ms. Akiko Sakai for technical assistance, and Mr. Takashi Arita for providing the experimental samples. This work was partially supported by a Grant-in-Aid for Scientific Research (18K05623) from the Japan Society for the Promotion of Science to Yukio Nagano. E.P.W. was supported by the Japanese Government (MEXT) scholarship. We would also like to thank Editage (https://www.editage.com) for the English language editing. This study was part of a dissertation submitted by the first author toward the partial fulfillment of her Ph.D. All authors provided consent for submission and publication.

## Author Contributions

E.P.W., Y.N., and T.I. designed the study. M.Y. and N.K. supplied the experimental samples. Y.N. performed the DNA experiments. E.P.W., T.I., M.H.S., and Y.N. conducted the bioinformatics analysis. N.U.J. established the framework for conducting the study, including an initial phase of data analysis conducted in Sri Lanka. As experts on Aurantioideae, M.Y. and N.K. reviewed the descriptions of Aurantioideae. All authors were involved in the data analysis. The manuscript was written by E.P.W. and Y.N. and reviewed by all authors.

## Data Availability Statement

Raw sequencing data have been deposited in the DDBJ Sequence Read Archive (https://www.ddbj.nig.ac.jp/dra/index-e.html; accession no. DRA017638). The cp genome sequences assembled and annotated from these raw sequencing data have been deposited in DDBJ/EMBL/GenBank under accession numbers LC794878 to LC794908. The alignments, tree data and Nexus files used in this study are included in the Supplementary Datasets 2 through 16.

## References

1. Tian, Y. et al. Research progress in plant molecular systematics of Lauraceae. Biology (Basel) 10, 391; 10.3390/biology10050391 (2021).

2. Jiang, D. et al. Complete chloroplast genomes provide insights into evolution and phylogeny of *Zingiber* (Zingiberaceae). BMC Genom. 24, 30; 10.1186/s12864-023-09115-9 (2023).

3. Li, C.C. et al. Insights into chloroplast genome evolution in Rutaceae through population genomics. Horticulture Advances 2, 13; 10.1007/s44281-024-00032-9 (2024).

4. Park, H.S. et al. Inheritance of chloroplast and mitochondrial genomes in cucumber revealed by four reciprocal F1 hybrid combinations. Sci. Rep. 11, 2506; 10.1038/s41598-021-81988-w (2021).

5. Sun, J. et al. Evolutionary and phylogenetic aspects of the chloroplast genome of *Chaenomeles* species. Sci. Rep. 10, 11466; 10.1038/s41598-020-67943-1 (2020).

6. Feng, J. et al. Analysis of complete chloroplast genome: Structure, phylogenetic relationships of *Galega orientalis* and evolutionary inference of Galegeae. Genes (Basel) 14, 176; 10.3390/genes14010176 (2023).

7. Kahraman, K. & Lucas, S.J., Comparison of different annotation tools for characterization of the complete chloroplast genome of *Corylus avellana* cv Tombul. BMC Genom. 20, 874; 10.1186/s12864-019-6253-5 (2019).

8. Wang, Y. et al. Chloroplast genome variation and phylogenetic relationships of *Atractylodes* species. BMC Genom. 22, 103; 10.1186/s12864-021-07394-8 (2021).

9. Greiner, S., Sobanski, J. & Bock, R., Why are most organelle genomes transmitted maternally? Bioessays 37, 80–94 (2015).

10. Rieseberg, L. H. & Soltis, D. E., Phylogenetic consequences of cytoplasmic gene flow in plants. Evol. Trend. Plant. 5, 65–84 (1991).

11. Pamilo, P. & Nei, M., Relationships between gene trees and species trees. Mol. Biol. Evol. 5, 568–583 (1988).

12. Soltis, D.E. & Kuzoff, R.K., Discordance between nuclear and chloroplast phylogenies in the Heuchera group (Saxifragaceae). Evolution 49, 727–742 (1995).

13. Thureborn, O., Wikström, N., Razafimandimbison, S.G. & Rydin, C., Plastid phylogenomics and cytonuclear discordance in Rubioideae Rubiaceae. PLoS One 19, e0302365; 10.1371/journal.pone.0302365 (2024).

14. Krawczyk, K., Nobis, M., Nowak, A., Szczecińska, M. & Sawicki, J., Phylogenetic implications of nuclear rRNA IGS variation in *Stipa* L. (Poaceae). Sci. Rep. 7,11506; 10.1038/s41598-017-11804-x (2017).

15. Wang, W., Li, H. & Chen, Z., Analysis of plastid and nuclear DNA data in plant phylogenetics—evaluation and improvement. Sci. China Life Sci. 57, 280–286 (2014).

16. Lutz, K.A., Wang, W., Zdepski, A. & Michael, T.P., Isolation and analysis of high quality nuclear DNA with reduced organellar DNA for plant genome sequencing and resequencing. BMC Biotechnol. 11, 54; 10.1186/1472-6750-11-54 (2011).

17. Amarasinghe, S.L. et al. Opportunities and challenges in long-read sequencing data analysis. Genome Biol. 21, 30; 10.1186/s13059-020-1935-5 (2020).

18. Mani, I., Current status and challenges of DNA sequencing. Advances in synthetic biology, 71– 80 (2020).

19. Wang, X., Xiong, T., Wang, Y., Zhang, X. & Sun, M., Integrating genomic sequencing resources: an innovative perspective on recycling with universal Angiosperms353 probe sets. Hortic. Adv. 2, 4; 10.1007/s44281-023-00026-z (2024).

20. Pezzini, F. F. et al. Target capture and genome skimming for plant diversity studies. APPS. 11, e11537; 10.1002/aps3.11537 (2023).

21. Dylus, D., Altenhoff, A., Majidian, S., Sedlazeck, F. J. & Dessimoz, C., Inference of phylogenetic trees directly from raw sequencing reads using Read2Tree. Nat. Biotechnol. 42, 139–147 (2024).

22. Cai, L., Zhang, H. & Davis, C.C., PhyloHerb: A highDthroughput phylogenomic pipeline for processing genome skimming data. APPS, 10, e11475; 10.1002/aps3.11475 (2022).

23. Andermann, T., Cano, Á., Zizka, A., Bacon, C. & Antonelli, A., SECAPR—a bioinformatics pipeline for the rapid and user-friendly processing of targeted enriched Illumina sequences, from raw reads to alignments. PeerJ, 6: e5175; 10.7717/peerj.5175 (2018).

24. Yang, F., Ge, J., Guo, Y., Olmstead, R. & Sun, W., Deciphering complex reticulate evolution of Asian Buddleja (Scrophulariaceae): insights into the taxonomy and speciation of polyploid taxa in the Sino-Himalayan region. Ann. Bot. 132, 15–28 (2023).

25. Park, S., Kwak, M. & Park, S., Complete organelle genomes of Korean fir, *Abies koreana* and phylogenomics of the gymnosperm genus *Abies* using nuclear and cytoplasmic DNA sequence data. Sci. Rep. 14, 7636; 10.1038/s41598-024-58253-x (2024).

26. Soltis, P. S., & Soltis, D. E., The role of hybridization in plant speciation, Annu. Rev. Plant Biol. 60, 561–588 (2009).

27. Rieseberg, L. H., & Willis, J. H., Plant speciation. Science 317, 910–914 (2007).

28. Nelson, T.C. et al. Ancient and recent introgression shape the evolutionary history of pollinator adaptation and speciation in a model monkeyflower radiation (*Mimulus* section *Erythranthe*). PLoS Genet. 17, e1009095; 10.1371/journal.pgen.1009095 (2021).

29. Suvorov, A. et al. Deep ancestral introgression shapes evolutionary history of dragonflies and damselflies. Syst. Biol. 71, 526–546 (2022).

30. Maddison, W. P., Gene trees in species trees. Syst. Biol. 46, 523–536 (1997).

31. Meleshko, O. et al. Extensive genome-wide phylogenetic discordance is due to incomplete lineage sorting and not ongoing introgression in a rapidly radiated *Bryophyte* genus. Mol Biol Evol. 38, 2750–2766 (2021).

32. Kandziora, M., Sklenář, P., Kolář, F. & Schmickl, R., How to tackle phylogenetic discordance in recent and rapidly radiating groups? Developing a workflow using *Loricaria* (Asteraceae) as an example. Front. Plant Sci. 12, 765719; 10.3389/fpls.2021.765719 (2022).

33. Wanke, S. & Wicke, S., Phylogenomic discordance in plant systematics. Front. Plant Sci. 14, 1308126; 10.3389/fpls.2023.1308126 (2023).

34. Nagano, Y. et al. Phylogenetic relationships of Aurantioideae (Rutaceae) based on RAD-Seq. Tree Genet. Genomes 14, 6; 10.1038/nrg.2015.28 (2018).

35. Herrero, R., Asins, M.J., Carbonell, E.A. & Navarro, L., Genetic diversity in the orange subfamily Aurantioideae. I. Intraspecies and intragenus genetic variability. Theor. Appl. Genet. 92, 599–609 (1996).

36. Penjor, T. et al. Phylogenetic relationships of *Citrus* and its relatives based on *matK* gene sequences. PLoS One 8, e62574; 10.1371/journal.pone.0062574 (2013).

37. Gabaldón, T. & Koonin, E.V., Functional and evolutionary implications of gene orthology. Nat. Rev. Genet. 14, 360–366 (2013).

38. Swingle, W. T. & Reece, P. C., The botany of Citrus and its wild relatives in the orange subfamily. In The citrus industry, vol 1 (eds. Reuther, W., Webber, H. J. & Bachelor, L. D.) 190–430 (University of California, Berkeley, 1967).

39. Malinsky, M., Matschiner, M. & Svardal, H., Dsuite-Fast D-statistics and related admixture evidence from VCF files. Mol. Ecol. Resour. 21, 584–595 (2021).

40. Pease, J.B. & Hahn, M.W., Detection and polarization of introgression in a five-taxon phylogeny. Syst. Biol. 64, 651–662 (2015).

41. Edelman, N.B. et al. Genomic architecture and introgression shape a butterfly radiation. Science 366, 594–599 (2019).

42. Hodel, R. G. J., Zimmer, E. A., Liu, B. B., & Wen, J., Synthesis of nuclear and chloroplast data combined with network analyses supports the polyploid origin of the apple tribe and the hybrid origin of the Maleae-Gillenieae Clade. Front. Plant Sci. 12, 820997; 10.3389/fpls.2021.820997 (2022).

43. Yu, W. B., Huang, P. H., Li, D. Z. & Wang, H., Incongruence between nuclear and chloroplast DNA phylogenies in *Pedicularis* section *Cyathophora* (Orobanchaceae). PLoS One 8, e74828; 10.1371/journal.pone.0074828 (2013).

44. Bruun-Lund, S., Clement, W. L., Kjellberg, F., & Rønsted, N., First plastid phylogenomic study reveals potential cyto-nuclear discordance in the evolutionary history of *Ficus* L. (Moraceae). Mol. Phylogenet. Evol. 109, 93–104 (2017).

45. Morales-Briones, D.F. et al. Disentangling sources of gene tree discordance in phylogenomic data sets: testing ancient hybridizations in Amaranthaceae s.l *Syst*. Biol. 70, 219–235 (2021).

46. Hibbins, M.S., Breithaupt, L.C. & Hahn, M.W., Phylogenomic comparative methods: Accurate evolutionary inferences in the presence of gene tree discordance. PNAS 120, e2220389120; 10.1073/pnas.2220389120 (2023).

47. Mendes, F.K., Hahn, Y. & Hahn, M.W., Gene tree discordance can generate patterns of diminishing convergence over time. Mol. Biol. Evol. 33, 3299–3307 (2016).

48. Bleidorn, C. & Bleidorn, C., Sources of error and incongruence in phylogenomic analyses. In Phylogenomics: an introduction, 173–193; 10.1007/978-3-319-54064-1_9 (2017). (2017).

49. Liu, X. et al. Genomic insights into zokors’ phylogeny and speciation in China. PNAS 119, e2121819119; 10.1073/pnas.2121819119 (2022).

50. Hernández-Gutiérrez, R. et al. Localized Phylogenetic Discordance Among Nuclear Loci Due to Incomplete Lineage Sorting and Introgression in the Family of Cotton and Cacao (Malvaceae). Front. Plant Sci. 13, 850521; 10.3389/fpls.2022.850521 (2022).

51. Yasuda, K. et al. Identification of parental chromosomes in sexual intergeneric hybrid progenies between *Citrus* cultivar ‘Nanpu’ tangor and *Citropsis schweinfurthii* in the subfamily Aurantioideae. J. Japan. Soc. Hort. Sci. 79, 129–134 (2010).

52. Smith, M.W., Gultzow, D.L. & Newman, T.K., First fruiting intergeneric hybrids between *Citrus* and *Citropsis*. J. Am. Soc. Hort. Sci. 138, 57–63 (2013).

53. Ramadugu, C. et al. A six nuclear gene phylogeny of *Citrus* (Rutaceae) taking into account hybridization and lineage sorting. PLoS One 8, e68410; 10.1371/journal.pone.0068410 (2013).

54. Fu, X.G. et al. Phylogenomic analysis of the hemp family (Cannabaceae) reveals deep cytoDnuclear discordance and provides new insights into generic relationships. J. Syst. Evol. 61, 806–826 (2023).

55. Aardema, M.L., Schmidt, K.L. & Amato, G., Patterns of cytonuclear discordance and divergence between subspecies of the scarlet macaw (*Ara macao*) in Central America. Genetica 151, 281–292 (2023).

56. Duan, L., Fu, L. & Chen, H.F., Phylogenomic cytonuclear discordance and evolutionary histories of plants and animals. Sci. China Life Sci. 66, 2946–2948 (2023).

57. Phuong, M.A., Bi, K. & Moritz, C., Range instability leads to cytonuclear discordance in a morphologically cryptic ground squirrel species complex. Mol. Ecol. 26, 4743–4755 (2017).

58. Hibbins, M.S. & Hahn, M.W., The timing and direction of introgression under the multispecies network coalescent. Genetics 211,1059–1073 (2019).

59. Seixas, F.A., Boursot, P. & Melo-Ferreira, J., The genomic impact of historical hybridization with massive mitochondrial DNA introgression. Genome Biol. 19, 91; 10.1186/s13059-018-1471-8 (2018).

60. Nge, F.J., Biffin, E., Thiele, K.R. & Waycott, M., Reticulate evolution, ancient chloroplast haplotypes, and rapid radiation of the Australian plant genus *Adenanthos* (Proteaceae). Front. Ecol. Evol. 8, 616741; 10.3389/fevo.2020.616741 (2021).

61. Chen, S., Zhou, Y., Chen, Y., & Gu, J., fastp: an ultra-fast all-in-one FASTQ preprocessor. Bioinform. 34, i884–i890 (2018).

62. Capella-Gutiérrez, S., Silla-Martínez, J. M. & Gabaldón, T., trimAl: A tool for automated alignment trimming in large-scale phylogenetic analyses. Bioinform. 25, 1972–1973 (2009).

63. Darriba, D. et al. ModelTest-NG: A new and scalable tool for the selection of DNA and protein evolutionary models. Mol. Biol. Evol. 37, 291–294 (2020).

64. Stamatakis, A., RAxML version 8: A tool for phylogenetic analysis and post-analysis of large phylogenies. Bioinform. 30, 1312–1313 (2014).

65. Huson, D. H. & Scornavacca, C., Dendroscope 3: An interactive tool for rooted phylogenetic trees and networks. Syst. Biol. 61, 1061–1067 (2012).

66. Jin, J. J. et al. GetOrganelle: A fast and versatile toolkit for accurate de novo assembly of organelle genomes. Genome Biol. 21, 241; 10.1186/s13059-020-02154-5 (2020).

67. Langmead, B. & Salzberg, S. L., Fast gapped-read alignment with Bowtie 2. Nat. Methods 9, 357–359 (2012).

68. Li, H. et al. The sequence alignment/map format and SAMtools. Bioinform. 25, 2078–2079 (2009).

69. Robinson, J. T., Thorvaldsdóttir, H., Turner, D. & Mesirov, J. P., igv. js: An embeddable JavaScript implementation of the Integrative Genomics Viewer (IGV). Bioinform. 39, btac830; 10.1093/bioinformatics/btac830 (2023).

70. Tillich, M. et al. GeSeq: versatile and accurate annotation of organelle genomes. Nucleic Acids Res. 45, W6–W11 (2017).

71. Zhong, X., Assembly, annotation and analysis of chloroplast genomes. Doctoral thesis in the University of Western Australia, Australia (2020).

72. Chan, P. P., Lin, B. Y., Mak, A. J. & Lowe, T. M., tRNAscan-SE 2.0: improved detection and functional classification of transfer RNA genes. Nucleic Acids Res. 49, 9077–9096 (2021).

73. Potter, S. C., et al. HMMER web server: 2018 update. Nucleic Acids Res. 46, W200–W204 (2018).

74. Greiner, S., Lehwark, P. & Bock, R., *OrganellarGenomeDRAW (OGDRAW)* version 1.3.1: Expanded toolkit for the graphical visualization of organellar genomes. Nucleic Acids Res. 47, W59–W64 (2019).

75. Bi, G., Mao, Y., Xing, Q. & Cao, M., HomBlocks: A multiple-alignment construction pipeline for organelle phylogenomics based on locally collinear block searching. Genomics 110, 18–22 (2018).

76. Drummond, A. J., Suchard, M. A., Xie, D. & Rambaut, A., Bayesian phylogenetics with BEAUti and the BEAST 1.7. Mol. Biol. Evol. 29, 1969–1973 (2012).

77. Zhang, C., Rabiee, M., Sayyari, E. & Mirarab, S., ASTRAL-III: polynomial time species tree reconstruction from partially resolved gene trees. BMC Bioinform. 19, 153; 10.1186/s12859-018-2129-y (2018).

78. Smith, S.A., Moore, M.J., Brown, J.W. & Yang, Y., Analysis of phylogenomic datasets reveals conflict, concordance, and gene duplications with examples from animals and plants. BMC Ecol. Evol. 15, 150; 10.1186/s12862-015-0423-0 (2015).

79. Wen, D., Yu, Y., Zhu, J. & Nakhleh, L., Inferring phylogenetic networks using PhyloNet. Syst. Biol. 67, 735–740 (2018).

80. Sullivan, J., & Joyce, P., Model selection in phylogenetics. Annu. Rev. Ecol. Evol. Syst. 36, 445–466 (2005).

81. Akaike, H., In Selected Papers of Hirotugu Akaike (eds Emanuel Parzen, Kunio Tanabe, & Genshiro Kitagawa) 199–213 (Springer New York, 1998).

