## Supplementary material for "Subfamily evolution analysis using nuclear and chloroplast data from the same reads": Legends for Supplementary Figures

### Supplementary Figure S1.

The ML phylogenetic tree was based on conserved nuclear gene sequences of 44 members of the Rutaceae family. The alignment used for the phylogenetic tree construction included 6,656,811 nucleotide positions, with 798,769 parsimony-informative sites, as calculated using custom Python scripts (Supplementary Dataset 1). RAxML analysis utilized the optimal evolutionary model GTRGAMMAIX, with *Ailanthus altissima* serving as the outgroup. Bootstrap values, represented as percentages of over 100 replicates, are displayed at the nodes, and the scale bar indicates genetic divergence in substitutions per site. Values below 50% are omitted. The subgroups identified in this study are distinguished by different colors. The alignment data in FASTA format and tree data in Newick format are available in Supplementary Dataset 7.

### Supplementary Figure S2.

The ML phylogenetic tree was constructed using the whole cp genome sequences of 59 species from the Rutaceae family. The alignment used for the phylogenetic tree construction included 59,231 nucleotide positions with 7,216 parsimony-informative sites, as calculated using MEGA11. The optimal GTRGAMMAIX evolutionary model was applied in the RAxML analysis, with *Ailanthus altissima* designated as the outgroup. Bootstrap values, derived from 1,000 replicates, are displayed at the nodes, excluding values below 50%. The scale bar illustrates genetic divergence in substitutions per site. Sequences assembled *de novo* are marked with asterisks. The subgroups identified in this study are differentiated by distinct colors. The alignment data in FASTA format and tree data in Newick format are available in Supplementary Dataset 8.

### Supplementary Figure S3.

The ML phylogenetic tree was constructed using the whole cp genome sequences of 39 species in the Aurantioideae subfamily.  The alignment used for the phylogenetic tree construction included a total of 88,197 nucleotide positions, with 4,215 parsimony-informative sites, as calculated using MEGA11. The GTRGAMMAIX model was chosen as the optimal evolutionary model and applied using RAxML. The root position of this tree aligns with that found in the phylogenetic tree of the Rutaceae family (Supplementary Fig. S2). Bootstrap values, presented as percentages over 1,000 replicates, are shown at the nodes. Bootstrap values falling below 50% are not included. The scale bar denotes genetic divergence in terms of substitutions per site. Asterisks highlight sequences assembled *de novo* in this study. Different colors denote the subgroups identified in this study. Since the DNA sequences of *Atalantia monophyla* and *Atalantia spinosa*, along with those of *Citropsis schweinfurthii* and *Citropsis gilletiana*, were completely identical, we did not include the data for *Atalantia spinosa* and *Citropsis gilletiana* in this study*.* The alignment data in FASTA format and the tree data in Newick format are available in Supplementary Dataset 3.

### Supplementary Figure S4.

The BI phylogenetic tree, constructed using the whole cp genome sequences of 39 species in the Aurantioideae subfamily, included a total of 88,197 nucleotide positions with 4,215 parsimony-informative sites. The BI support values are indicated at each node. The root position of this tree aligns with that found in the phylogenetic tree of the Rutaceae family (Supplementary Fig. S2). The scale bar denotes genetic divergence in terms of substitutions per site. Asterisks highlight sequences assembled *de novo* in this study. The subgroups identified in this study are differentiated by distinct colors. The alignment data in FASTA format and tree data in Newick format are available in Supplementary Dataset 9.

### Supplementary Figure S5.

Discordance Between Concatenated Nuclear Gene Trees and ASTRAL Species Tree

(A) Comparison of the concatenated nuclear gene tree derived from Read2Tree and the concatenated nuclear gene tree based on the subset of 579 genes. Left panel: ML phylogenetic tree generated from conserved nuclear gene sequences of 39 species within the Aurantioideae subfamily using Read2Tree. Bootstrap support values (percentages from 100 replicates) are indicated at each node, with values below 50% omitted. Subgroups identified in this study are depicted using distinct colors.

(B) Comparison of the ASTRAL species tree with the concatenated nuclear gene tree based on the 579-gene subset. Left panel: Species tree inferred using ASTRAL-III, with local posterior probability (LPP) values shown as percentages for each node. Subgroups identified in this study are highlighted using different colors. The alignment of the selected 579 genes is available in Supplementary Dataset 10, and tree data in Newick format can be found in Supplementary Dataset 4.

Right panel in both (A) and (B): ML concatenated nuclear gene tree constructed from the 579-gene subset. Bootstrap support values (percentages from 100 replicates) are displayed at the nodes, with values below 50% omitted. Subgroups are represented with distinct colors. Dotted lines and shapes emphasize regions of conflicting relationships, illustrating topological incongruence between the two trees. Alignment data in FASTA format and tree data in Newick format are available in Supplementary Dataset 11.

### Supplementary Figure S6.

Phylogenetic networks for Citrus and its near groups were inferred using PhyloNet with the Maximum Pseudolikelihood method under one, two, and three reticulation models. Green branches denote lineages associated with reticulate evolutionary histories, and numerical values indicate the inheritance probabilities for each reticulation event.

### Supplementary Figure S7.

Phylogenetic networks for the Bael fruit group and its associated species were inferred using PhyloNet with the Maximum Pseudolikelihood method under one, two, and three reticulation models. Purple branches denote lineages associated with reticulate evolutionary histories, and numerical values indicate the inheritance probabilities for each reticulation event.
