## Supplementary figures and images for "Subfamily evolution analysis using nuclear and chloroplast data from the same reads"

### Supplementary Fig S1

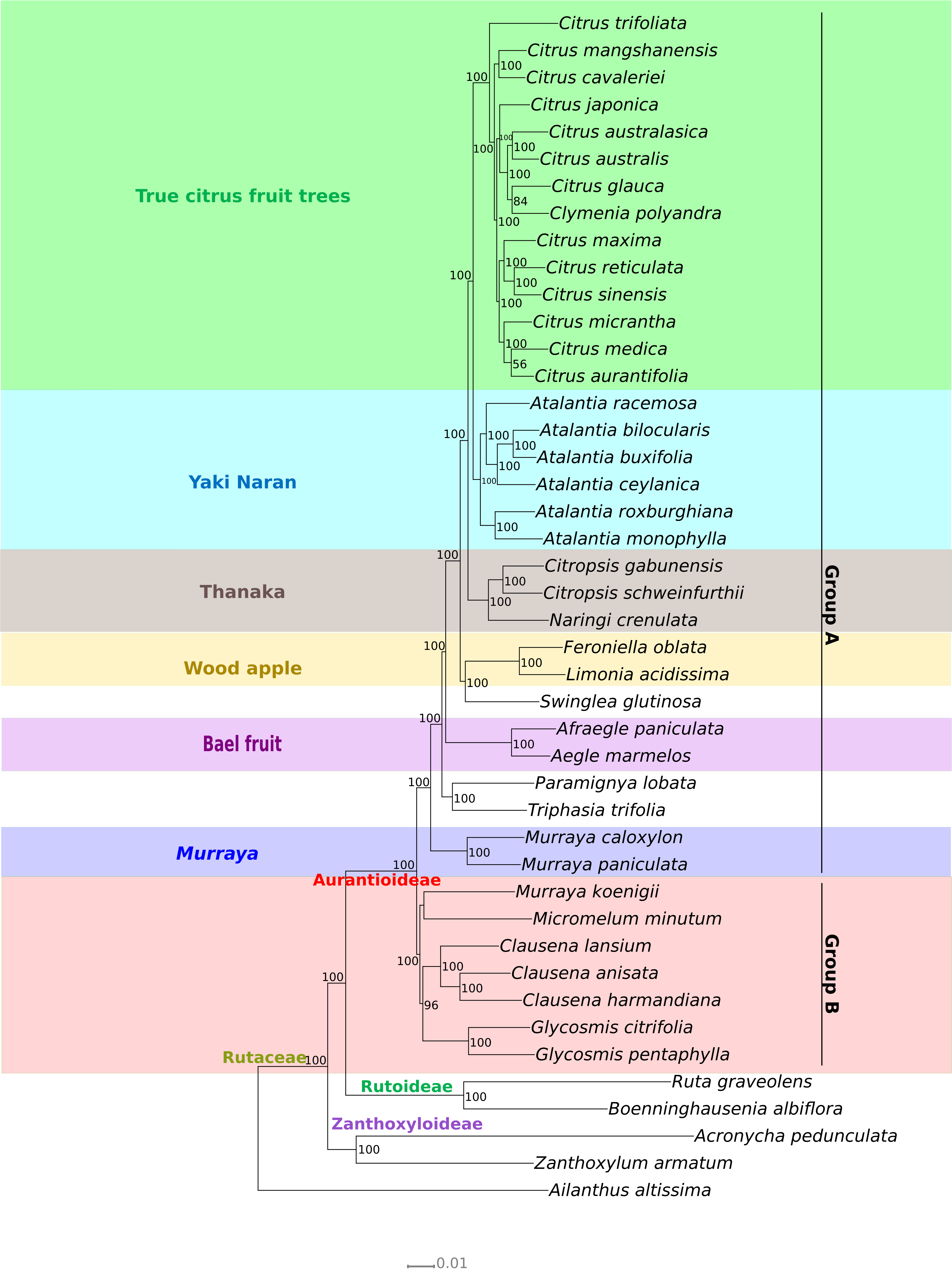

### Supplementary Fig S2

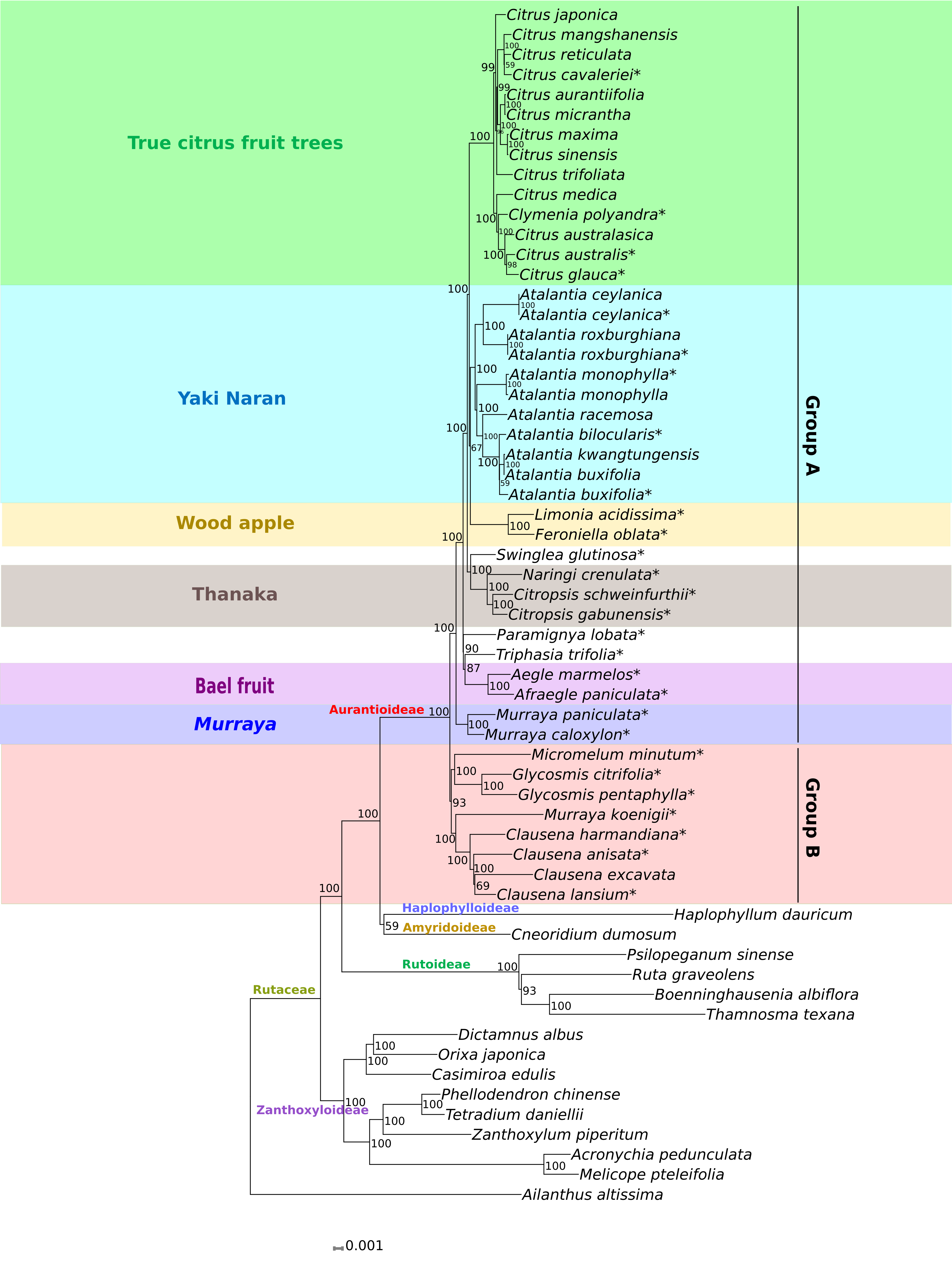

### Supplementary Fig S3

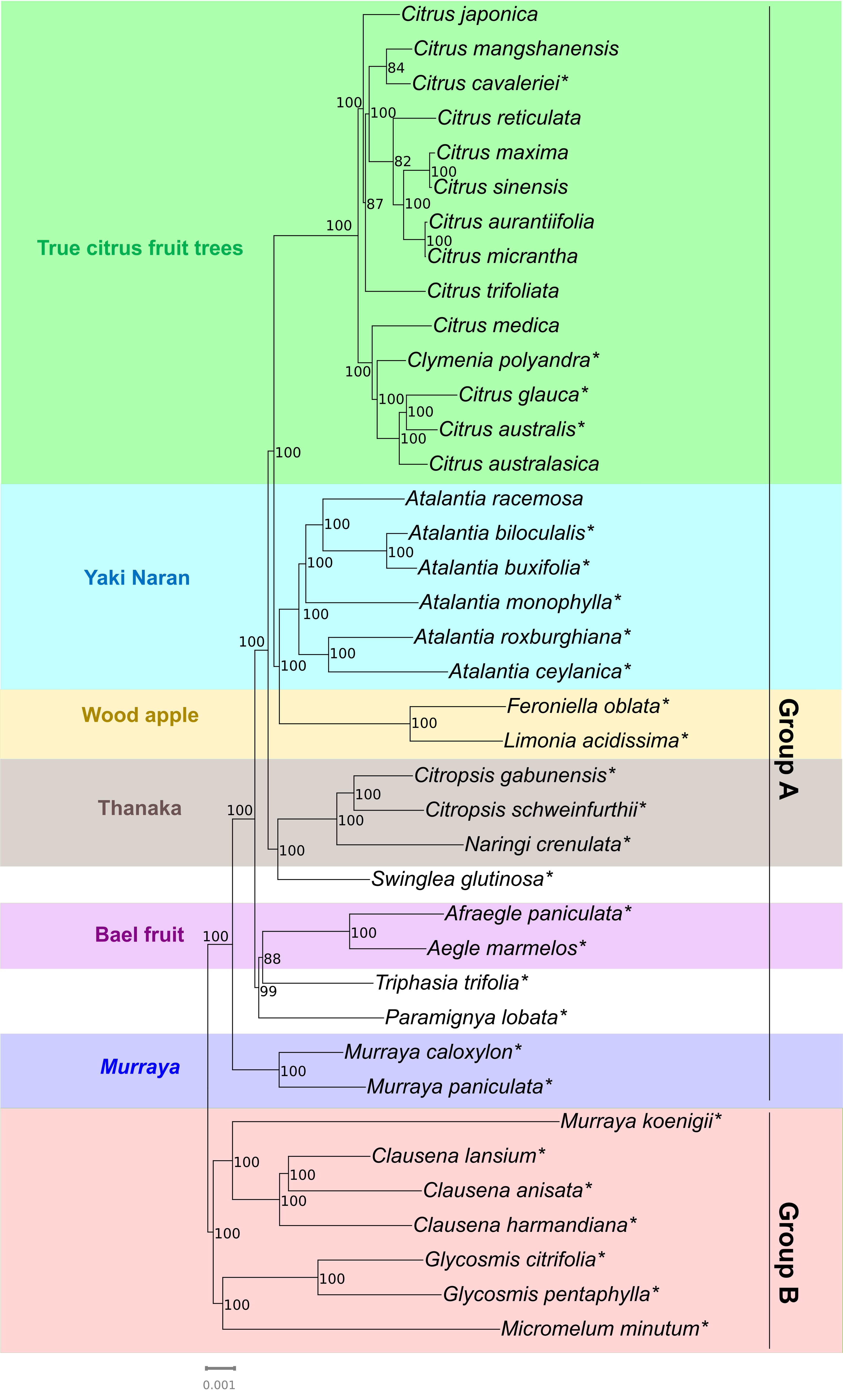

### Supplementary Fig S4

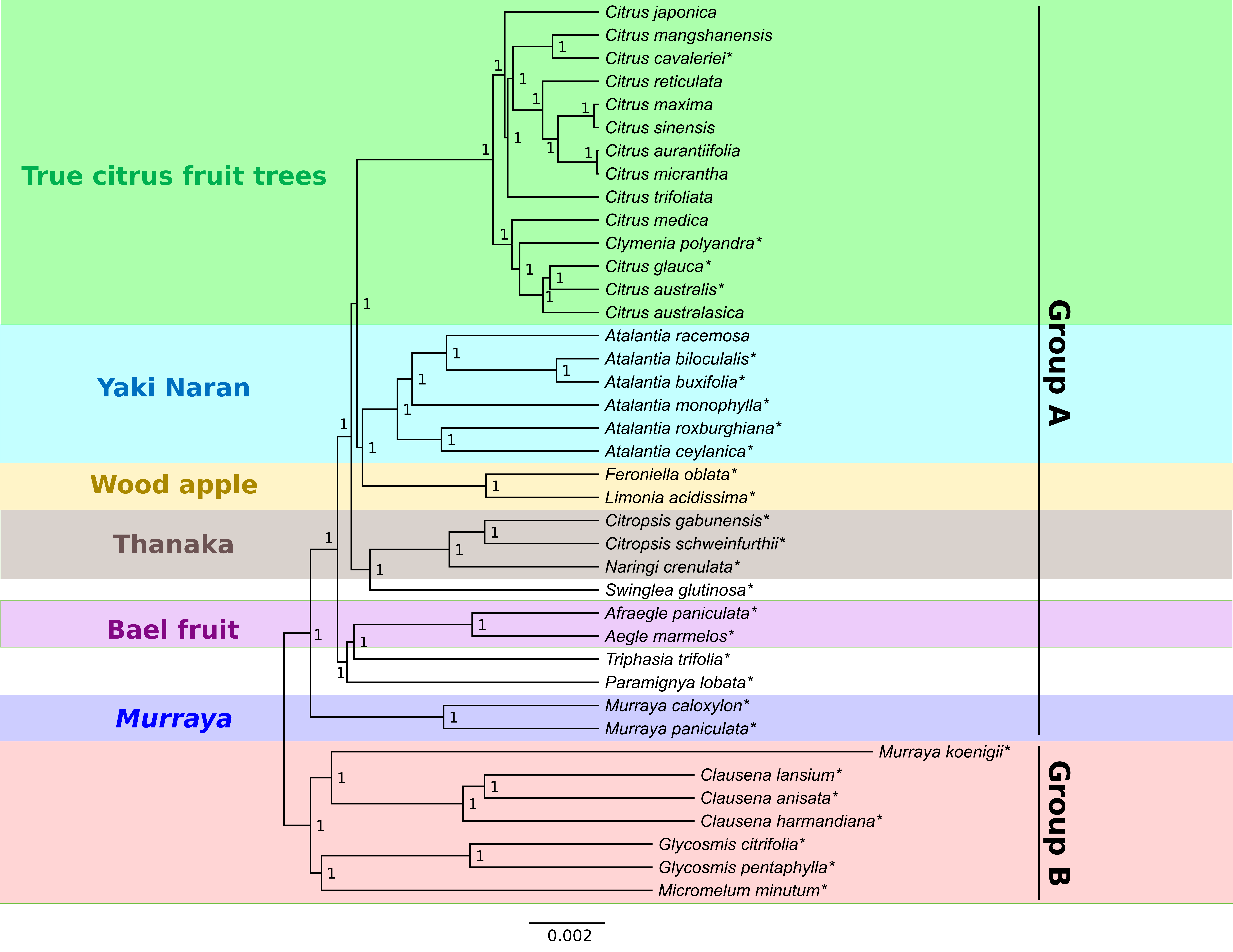

### Supplementary Fig S5

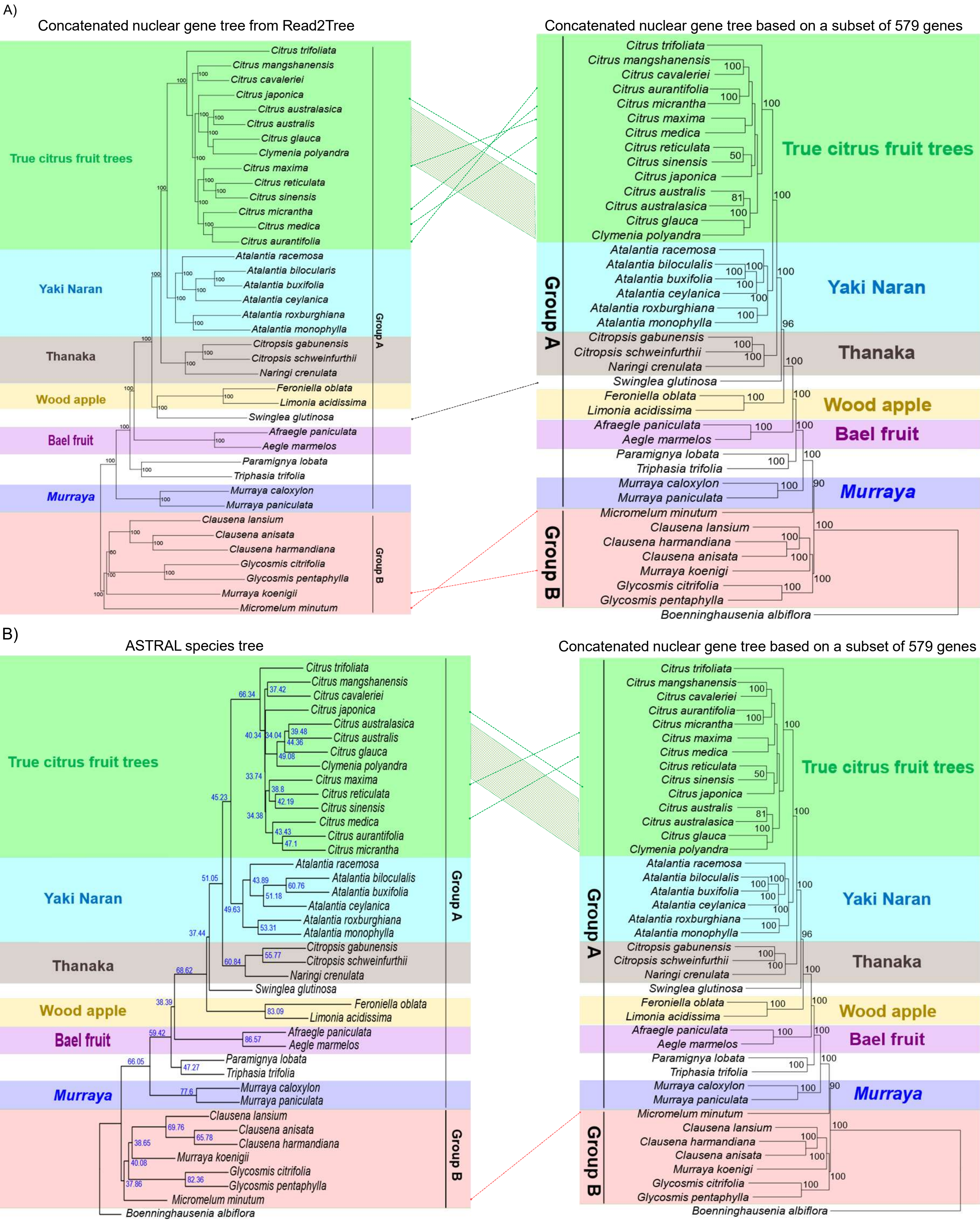

### Supplementary Fig S6

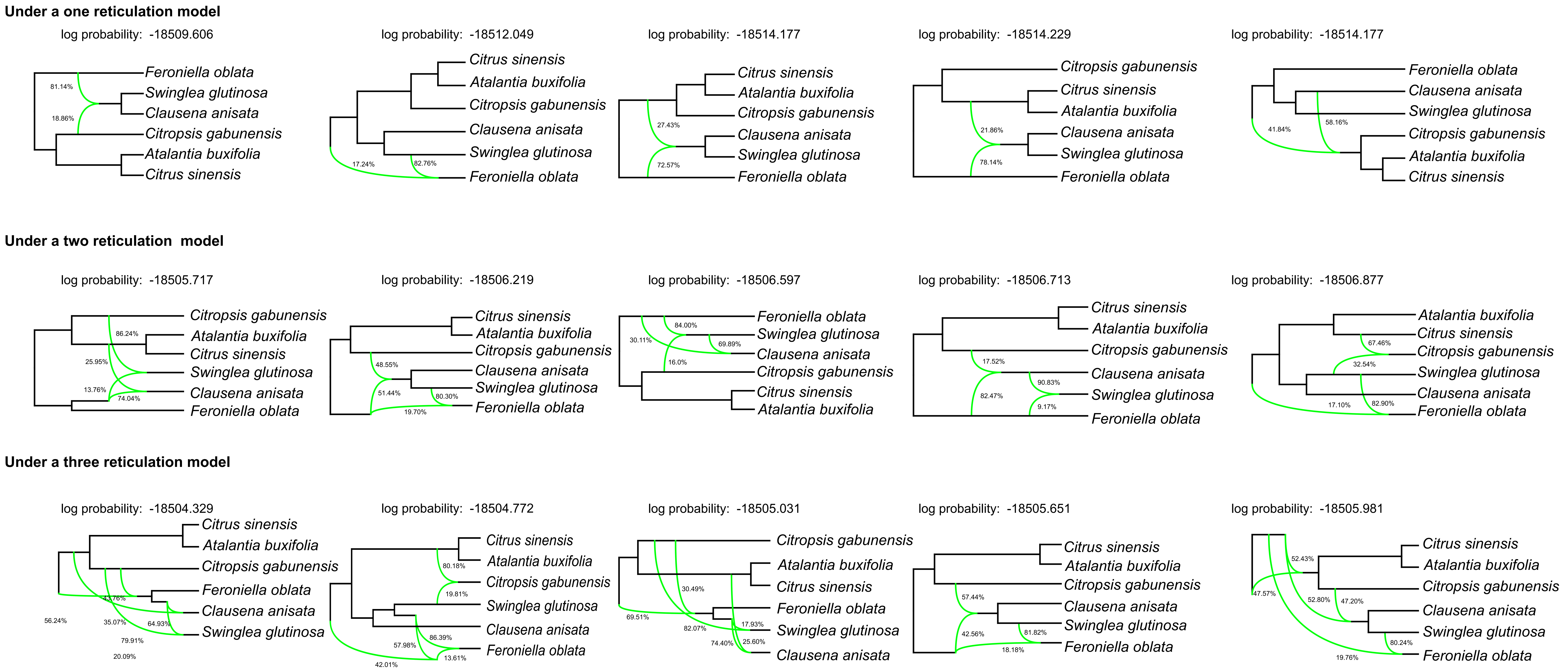
