## Supplementary Fig S7 for "Subfamily evolution analysis using nuclear and chloroplast data from the same reads"

### Under a one reticulation model

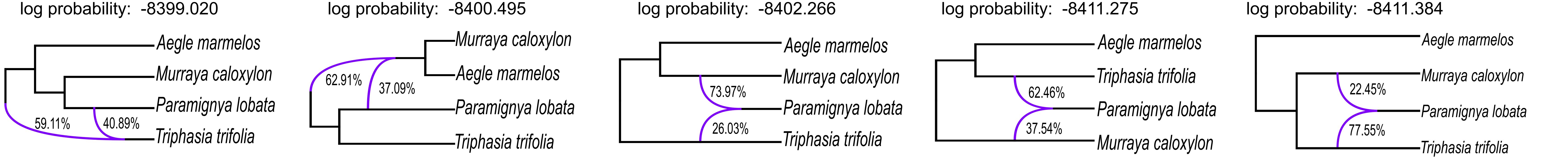


### Under a two reticulation model

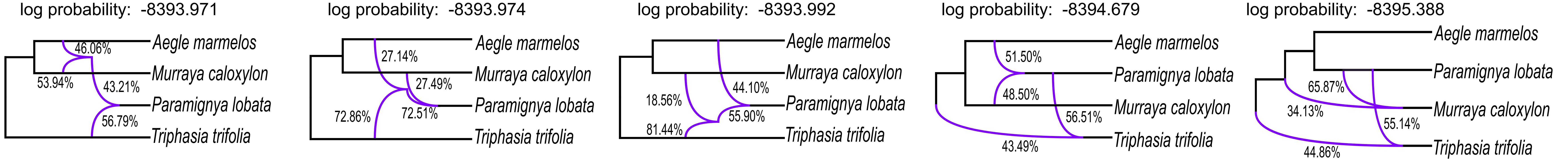


### Under a three reticulation model

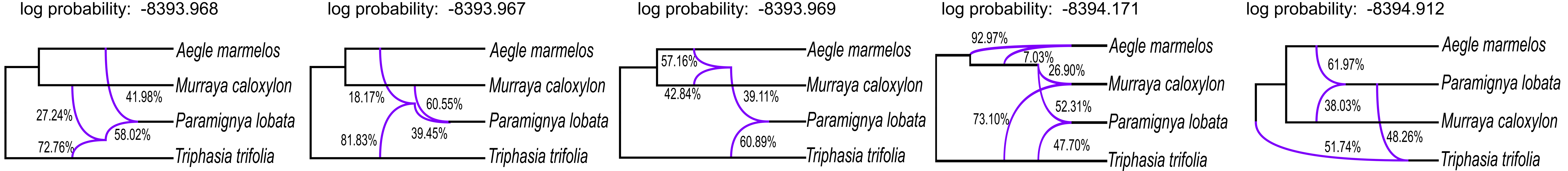
