## Supplementary Dataset 1 for "Subfamily evolution analysis using nuclear and chloroplast data from the same reads": Custom_Python_scripts.txt

#Python script to analyze multiple sequence alignments and determine the length of each sequence in the alignment,

from Bio import AlignIO

# Load your multiple sequence alignment file
alignment = AlignIO.read("your_alignment_file.fasta", "fasta")

# Print the length of each sequence in the alignment
for record in alignment:
    print(f"Sequence {record.id}: Length {len(record.seq)}")

#Python script to analyze multiple sequence alignments and count the number of parsimony-informative sites, while treating "N" or "?" characters as missing data,

from Bio import AlignIO

def count_parsimony_informative_sites(alignment):
    informative_sites = 0
    for i in range(alignment.get_alignment_length()):
        column = alignment[:, i]
        # Filter out columns with missing data represented as 'N' or '?'
        filtered_column = [char for char in column if char not in ['N', '?']]
        unique_chars = set(filtered_column)
        # Count as informative only if two or more types each appear at least twice
        if len(unique_chars) > 1 and all(filtered_column.count(char) > 1 for char in unique_chars):
            informative_sites += 1
    return informative_sites

# Load your multiple sequence alignment file
alignment = AlignIO.read("your_alignment_file.fasta", "fasta")

# Count the parsimony-informative sites
informative_sites_count = count_parsimony_informative_sites(alignment)
print(f"Parsimony-Informative Sites: {informative_sites_count}")
